# Architecture of the AP2:clathrin coat on the membranes of clathrin-coated vesicles

**DOI:** 10.1101/2020.01.28.922591

**Authors:** Oleksiy Kovtun, Veronica Kane Dickson, Bernard T. Kelly, David. J. Owen, John A. G. Briggs

**Author notes:** correspondence to BTK, DJO or JAGB.

## Abstract

Clathrin-mediated endocytosis (CME) is crucial for modulating the protein composition of a cell’s plasma membrane. Clathrin forms a cage-like, polyhedral outer scaffold around a vesicle, to which cargo-selecting clathrin adaptors are attached. AP2 is the key adaptor in CME. Crystallography has shown AP2 to adopt a range of conformations. Here we used cryo-electron microscopy, tomography and subtomogram averaging to determine structures, interactions and arrangements of clathrin and AP2 at the key steps of coat assembly, from AP2 in solution to membrane-assembled clathrin-coated vesicles (CCVs). AP2 binds cargo and PtdIns(4,5)*P*_2_-containing membranes via multiple interfaces, undergoing conformational rearrangement from its cytosolic state. The binding mode of AP2 β2-appendage into the clathrin lattice in CCVs and buds implies how the adaptor structurally modulates coat curvature and coat disassembly.

## Introduction

CME is a vital multistep process in all eukaryotes in which plasma membrane cargo proteins are first concentrated into a patch by clathrin adaptors, which are themselves clustered by interaction with a polymeric clathrin scaffold. The patch of membrane is deformed towards the cytoplasm and undergoes scission from the parent membrane to form a clathrin-coated vesicle (CCV), which subsequently delivers its cargo into the endocytic system (reviewed in Kirchhausen et al., 2014; Mettlen et al., 2018; Mettlen and Danuser, 2014; Schmid et al., 2014; Taylor et al., 2011).

The clathrin lattice, containing both pentagonal and hexagonal openings, is formed from triskelia containing three copies of the clathrin heavy chain (CHC) and three copies of the light chain (CLC) (reviewed in Kirchhausen et al., 2014) (**Fig. S1**). The CHC N-terminal domain (NTD) β-propeller contains multiple binding sites for clathrin box peptides, which are scattered throughout the unstructured regions of clathrin adaptors (Muenzner et al., 2017; Willox and Royle, 2012). The most abundant clathrin adaptor (Borner et al., 2012) is the 300kDa heterotetrameric AP2 complex, consisting of α, β2, µ2, and σ2 subunits (**Fig. S1**). AP2 plays a central role in CME through binding plasma membrane PtdIns(4,5)*P*_2_, the two most common cargo motifs (YxxΦ and [ED]xxxL[LI]), clathrin, and other regulatory/accessory proteins (reviewed in Kirchhausen et al., 2014).

AP2 is one of the first proteins to arrive at a forming CCV (Taylor et al., 2011) and is also important in initiating clathrin polymerisation (Cocucci et al., 2012; Motley et al., 2003). The favoured although unproven model for AP2 and clathrin in CME suggests that cytosolic AP2 is closed and unable to bind cargo and that membrane-bound AP2 adopts a more open conformation in which cargo binding sites are accessible (Hollopeter et al., 2014; Jackson et al., 2010). A closed conformation has been seen under non-physiological conditions in crystals (Collins et al., 2002). A number of different possible structures for membrane-bound, cargo-binding ‘open’ forms of AP2 complexes have been determined in the absence of membrane (Jackson et al., 2010; Wrobel et al., 2019). Similarly, the highest-resolution structures of clathrin have been determined in the absence of a membrane and/or a characterised complement of folded adaptors. It therefore remains unclear how AP2 and clathrin interact with one another and the membrane to form CCVs. Here we present the structures, and relative arrangements of AP2 and clathrin under physiological buffer conditions and in membrane-associated vesicle coats.

## Results and Discussion

### AP2 structures

We assessed the conformation(s) adopted by recombinant AP2 cores in solution under physiological buffer conditions using single-particle cryo-electron microscopy (**Fig. S2A**). The bulk of AP2 (∼80%) adopted a closed conformation, the structure of which we determined to 3.8 Å. The remaining particles (∼20%) did not refine to a reliable structure (i.e. no alternative ‘open’ conformer was identified). The structure is similar to that seen under high-salt, high-pH conditions in the crystal structure containing inositol hexakis-phosphate (Collins et al., 2002): the C-terminal domain of µ2 (Cµ2) is located in the spatially complementary bowl created by the other subunits, while the YxxΦ and [ED]xxxL[LI] cargo motif binding sites are blocked by portions of the β2 subunit (**Fig. 1A**, **S3**) (Jackson et al., 2010).

**Fig. 1.**
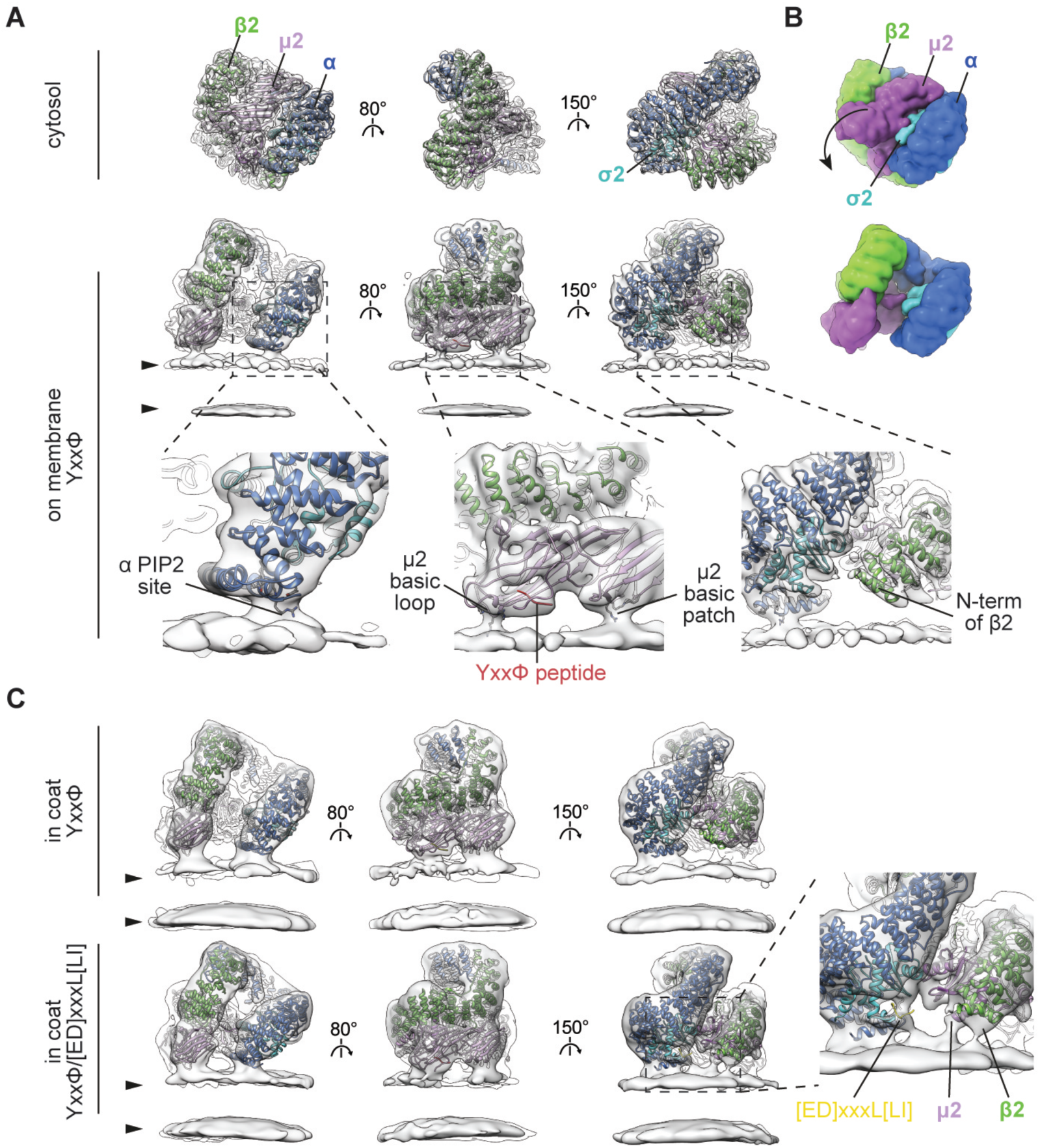
Conformational activation and membrane interactions of AP2. **A.** Comparison of single-particle structure of the cytosolic closed form (top row) with the tomography-resolved open form of AP2 on the membrane in the presence of YxxΦ cargo (lower rows). EM maps (semi-transparent grey) are fitted with corresponding ribbon models of AP2 colour-coded by chain. Arrowheads indicate membrane leaflets. Close-ups show details of membrane contacts with α and µ2 subunits and the lack of contact with β2, with PtdIns(4,5)*P_2_* sites and cargo peptides marked. **B.** Surfaces of the atomic models for cytosolic and membrane-recruited AP2 in **A** are shown to illustrate the rearrangement between cytosolic and membrane-bound forms. The arrow indicates the major movement of Cµ2 out of the bowl of AP2. **C.** Conformation and membrane interactions of AP2 recruited to the membrane via YxxΦ or YxxΦ/[ED]xxxL[LI] cargo signals within assembled clathrin coats. Additional tilting of α is apparent when [ED]xxxL[LI] cargo is present. The close-up demonstrates new membrane contacts formed by β2 and µ2 subunits, and additional density in the cargo binding pocket when [ED]xxxL[LI] cargo is present.

Recombinant AP2 FLAP (AP2 lacking the hinge and appendage of α; **Fig. S1**) (Kelly et al., 2014) was recruited onto phospholipid membranes containing PtdIns(4,5)*P*_2_ and TGN38 YxxΦ–cargo in the same physiological buffer conditions and imaged by cryo-electron tomography. AP2 coated existing liposomes but did not generate small buds or vesicles - AP2 alone does not drive membrane curvature/budding (**Fig. S2B**). We applied subtomogram averaging to determine the structure of AP2 to ∼9 Å resolution (**Fig. 1A**, **S3**). The structure is most similar to the previously-determined crystal structure of the open conformation (Jackson et al., 2010), although Cµ2 and the N-terminal portion of β2 are positioned approximately 5 Å further away from α (**Fig. S3C**). Compared to the closed form, Cµ2 has moved to form a planar membrane-binding surface (**Fig. 1B**), and its movement with respect to the β2 N-terminus has freed up the YxxΦ-cargo binding pocket, which has density consistent with bound cargo peptide. The [ED]xxxL[LI] cargo motif binding site on σ2 is accessible to membrane-embedded cargo due to a movement of the N-terminus of β2 relative to σ2. In the open form crystal structure (Jackson et al., 2010), the [ED]xxxL[LI] site is occupied by the Myc-tag of a neighbouring AP2. Our structure of AP2 on the membrane shows that the open conformation did not result from this or other crystal packing artefacts.

We observed three sites at which the AP2 electron density contacts the membrane: the proposed PtdIns(4,5)*P*_2_ binding site on the α subunit whose mutation abolishes AP2 membrane binding (Gaidarov et al., 1996; Honing et al., 2005; Jackson et al., 2010) and two putative PtdIns(4,5)*P*_2_ binding sites on Cµ2 (**Fig. 1A**), simultaneous mutation of which inhibits membrane binding and reduces CCV nucleation (Jackson et al., 2010; Kadlecova et al., 2017). The first Cµ2 site corresponds to a basic loop including K167, Y168, R169, R170 following the µ2 linker helix, which undergoes an extended to helical transition between closed and open AP2 (Jackson et al., 2010). This location suggests a direct physical connection between membrane binding and stabilisation of the open AP2 conformation and points to interference with this linkage being the explanation for why an R170W mutation impairs CME and causes neurological defects (Helbig et al., 2019). The second Cµ2 contact appears to be mediated by residues K350, K367, R368 within a basic patch also containing K330, K334, K352, K354, K356 and K365. We do not observe a membrane contact at the proposed β2 N-terminal PtdIns(4,5)*P*_2_-binding site (Honing et al., 2005; Jackson et al., 2010; Kadlecova et al., 2017).

We assembled both AP2 and clathrin onto YxxΦ cargo/PtdIns(4,5)*P*_2_-containing membranes. In contrast to the sample lacking clathrin, buds and vesicles were now formed with both AP2 and clathrin coating their membranes (**Fig. S2**) indicating a role for clathrin scaffold recruitment in membrane curvature. The AP2 structure was identical up to the determined resolution (12 Å) to that seen in the absence of clathrin (**Fig. 1C**). Inclusion of an additional ‘dileucine’ [ED]xxxL[LI]-containing cargo resulted in tilting of α, and movement of β2 away from σ2 to further expose its [ED]xxxL[LI] binding site. Helices 2 and 5 in σ2 are poorly resolved, and the proposed β2 N-terminal PdIns(4,5)*P_2_* binding site now contacts the membrane as does a region in the vicinity of helices 2 and 3 of µ2 (**Fig. 1C**).

When combined with published information, our data suggest a model whereby AP2 initially contacts the membrane in the closed conformation via PtdIns(4,5)*P*_2_ binding sites on α and β2. Liganding of the two Cµ2 PtdIns(4,5)*P_2_*-binding sites will shift the equilibrium to the ‘open’ state in which the cargo binding sites become unblocked. *In vivo* this may be aided by the binding of FCHO1/FCHO2/sgip (Hollopeter et al., 2014; Umasankar et al., 2014). AP2 can then ‘scan’ the local membrane for cargo and its binding would then further stabilise an open form on the membrane. The binding of PtdIns(4,5)*P*_2_ and cargo are thus allosterically linked.

The transitions from cytosolic to single-cargo membrane-bound AP2, and from single-to double-cargo membrane-bound AP2, involve conformational flexing of the α and β2 solenoids. We did not observe previously described open+ (Wrobel et al., 2019), AP1-like hyper-open (Jia et al., 2014) or splayed, COPI-like (Dodonova et al., 2015) forms (**Fig. S3**), suggesting that the open form is the lowest energy conformer for membrane associated, non-phosphorylated AP2.

### AP2 distribution on membranes

We found the distribution of AP2 on the surface of membranes to be largely irregular, lacking long-range organization. On YxxΦ-only containing membranes, ∼20% of AP2s are dimerised (**Fig. S4**), their β2 N-terminal helices having apparently moved to participate in the interface (**Fig. S4B**). The dimerisation interface is not similar to those proposed for AP1 multimerisation via the small GTPase Arf1 (Morris et al., 2018). The AP2 distribution was unchanged by addition of clathrin and hence by the formation of buds and vesicles. In vesicles and buds with both [ED]xxxL[LI] and YxxΦ cargoes, the dimeric form of AP2 is absent, likely because the N-terminal helix of β2 has moved to contact the membrane and can no longer contribute to dimerization interface. It has been speculated on the basis of crystal packing and single-particle electron microscopy (Ren et al., 2013; Shen et al., 2015) that AP1 and by analogy AP2 would adopt a regular lattice that would be stabilised/triggered by the presence of clathrin. There is, however, no AP2 lattice present in our assembled vesicles, excluding AP2 oligomerisation as a factor in membrane curvature generation.

### Clathrin structure

We determined the structure of clathrin above AP2 in both the YxxΦ and YxxΦ/[ED]xxxL[LI] cargo-containing liposomes by subtomogram averaging: the structures were the same (Materials and Methods), so were combined for further analysis. The clathrin legs were sorted into two classes according to whether the leg bounds a hexagonal or pentagonal cage face (**Fig. S5A,B**). Each class was averaged separately to generate structures at ∼7.7 Å resolution (**Fig. 2**, **S5C,D**). The CLC region that contacts CHC is well resolved, as are most of the leg and the helical hub of CHC. The resolution is lower towards the NTDs of CHC that extend downwards towards the membrane, reflecting mobility of these domains, but is sufficient to position the full NTD. Our structures are generally consistent with those previously described from purified clathrin in solution in the absence of membranes (Fotin et al., 2004b; Morris et al., 2019), but by now resolving the NTDs we were able to derive the positions of peptide binding within the clathrin cage (**Fig. 4, S5C**).

**Fig. 2.**
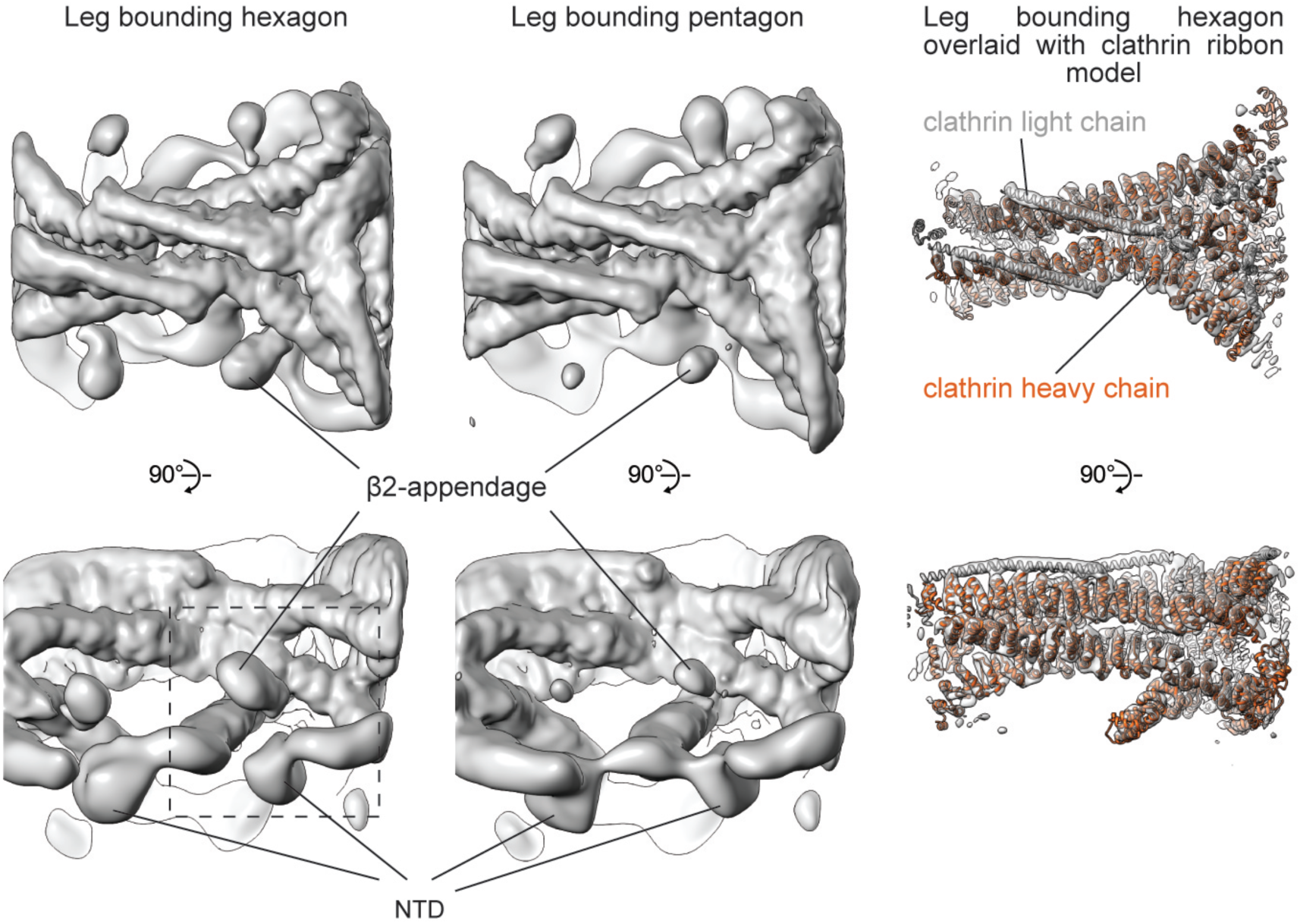
The structure of clathrin on coated membranes. EM maps (grey surface) of clathrin legs bounding hexagon and pentagon (see **Fig. S5** for details). The right column shows the EM map (sharpened to reveal high-resolution features) overlayed with a fitted ribbon model for the clathrin hub. N-terminal portions of CHC are clearly defined. Density corresponding to the AP2 β2-appendage indicates higher occupancy in hexagon than in pentagon maps (see also **Fig. S7**). The dashed box indicates region shown in **Fig. 4**.

### Clathrin/AP2 relationship

In YxxΦ cargo-containing coated buds, the clathrin NTDs are positioned at a higher radius than AP2, and there are no preferential positions or orientations of AP2 relative to clathrin (**Fig. 3**, **S6**). Inclusion of [ED]xxxL[LI] cargo, or transition of the buds to vesicles, both cause clathrin to move towards the membrane by 10-15 Å, after which the NTDs are at a lower radius than the top of AP2 (**Fig. S6**). In these cases AP2 is preferentially located in the gaps between the NTDs, though with no preferred rotational orientation. This suggests that the preferred localization is due to steric clashing rather than specific interaction. It is not clear to us what causes the movement of clathrin towards the membrane.

**Fig. 3.**
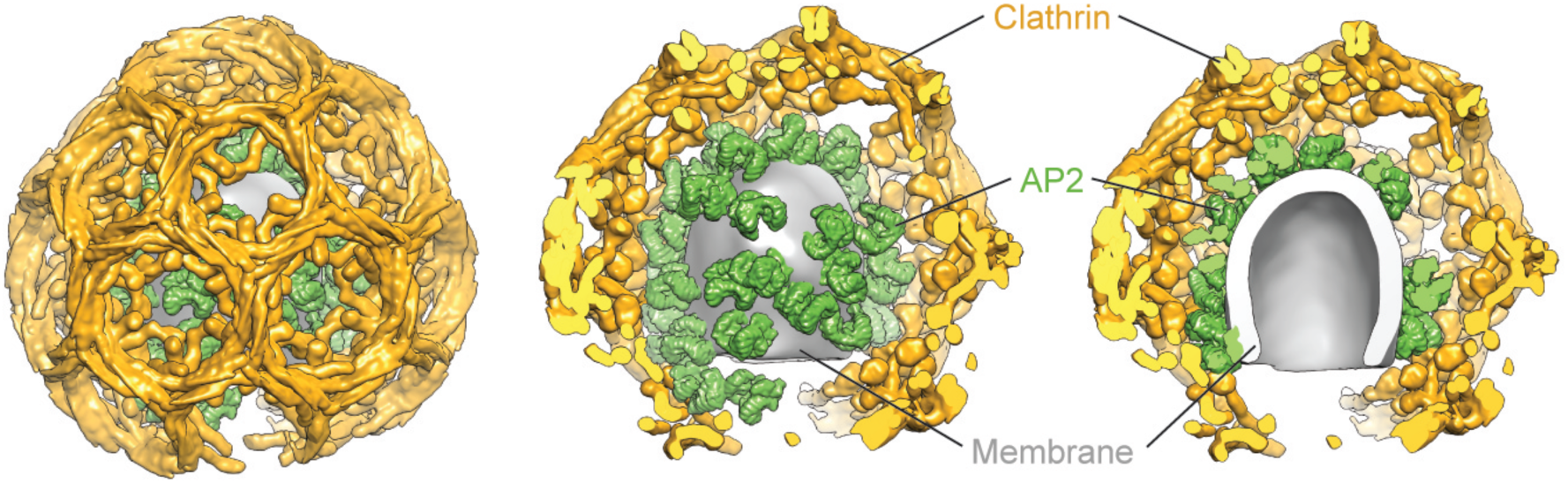
Experimental model of a coated bud formed on YxxΦ cargo-containing membrane. The model of a representative bud was produced by placing densities for AP2 (green) and clathrin legs (gold) at positions and orientations determined by subtomogram averaging for that bud. AP2 and clathrin (together with β2-appendage) densities were simulated on corresponding pseudoatomic models. The membrane was defined by segmenting the tomogram. The **left panel** shows the exterior of the bud. The front half of the clathrin cage is removed in the **middle panel**, revealing the random AP2 distribution on the membrane. The **right panel** shows a cross-section through the entire bud.

### AP2 β2-appendage

Between the clathrin NTDs and the undersides of the triskelia, we observed a ‘bean-shaped’ density that is not part of clathrin (**Fig. 2**). We applied principal component analysis to sort the datasets according to the presence or absence of this additional density (**Fig. S7A**), and averaged positions where it was present within a hexagon, obtaining a structure corresponding to the AP2 β2-appendage (Materials and Methods) (**Fig. 4A**, **S7**). The β2-hinge is not visible at this resolution. When comparing the density and occupancy of the β2-appendage between sites oriented towards the centres of either hexagons or pentagons, we found preferential binding in hexagonal faces (more than double the apparent occupancy at pentagonal faces, **Fig. S7A**).

**Fig. 4.**
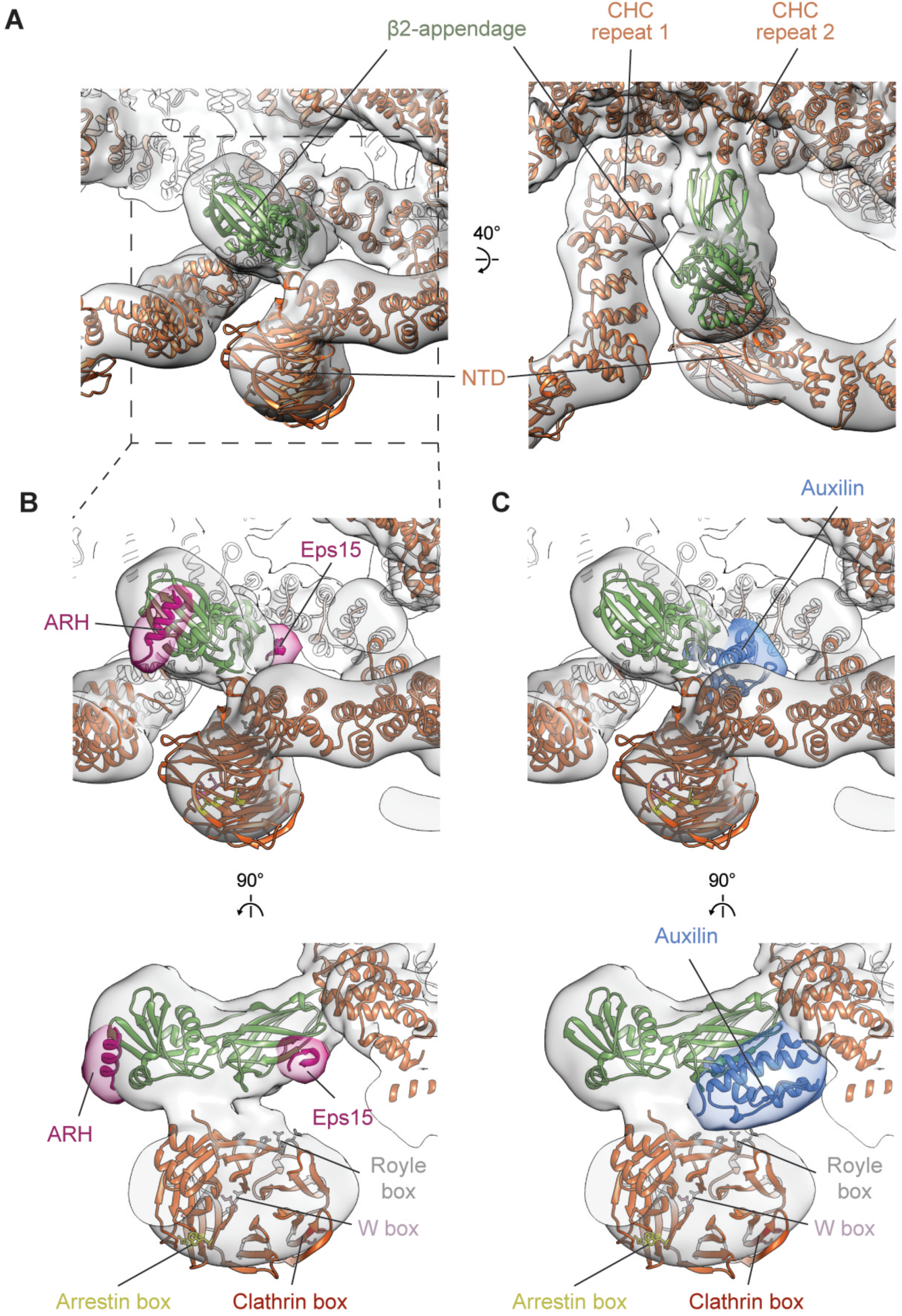
Interactions between clathrin and the AP2 β2-appendage domain. **A.** Positions of NTD and the β2-appendage with respect to the clathrin lattice. Overlay of EM map of leg bounding hexagon enriched in β2-appendage (transparent grey, see **Fig. S7** for details) with fitted protein ribbon models. The β2-appendage makes contact with sites on three CHC: two in the ankle segment (CHC repeats 1 and 2) and one in the NTD. **B.** Close-up of NTD and β2-appendage region indicating the binding sites of Low Density Lipoprotein Receptor adaptor ARH and Eps15. ARH and Eps15 peptides are pink, positioned as in PDB 2g30 (Edeling et al., 2006) and 2iv9 (Schmid et al., 2006), together with a simulated 15 Å resolution isosurface (transparent pink). Binding sites for clathrin boxes in the NTD are marked. **C.** As in B. showing the binding position of DNAJ Auxilin domain in blue, as described in PDB 1xi5 (Fotin et al., 2004a).

β2-appendages link CHC repeat 1 in one clathrin molecule, CHC repeat 2 in a second clathrin molecule (both in the ankle region), and the NTD of a third clathrin molecule (**Fig. 4A**). This positions the β2-appendage such that its C-terminal platform subdomain is freely accessible to the Fxx[FL]xxxR helical motifs on the unstructured portions of secondary adaptors such as ARH, arrestins and epsins (Edeling et al., 2006) (**Fig. 4B**) rather than being blocked as speculated (Schmid et al., 2006). Another motif-binding site, recognised by regulatory/accessory proteins including Eps15, is located on the β2-appendage sandwich domain close to the point of interaction with CHC repeat 2 (**Fig. 4B**).

The β2-appendage is dispensable for clathrin polymerisation in vitro (**Fig. S8**). Binding of the β2-appendage to the clathrin scaffold therefore likely reflects roles in regulating membrane curvature, assembly and/or disassembly in vivo. The β2-appendage clathrin binding site we can now see is directly adjacent to the proposed binding site of the DNAJ domain of auxilin (**Fig. 4C**). Auxilin can induce coat disassembly by recruiting HSC70 (Smith et al., 2004; Xing et al., 2010), which also could indirectly modulate levels of cargo incorporation by AP2 (Chen et al., 2019). We now see that auxilin would occupy a cavity bounded by the β2-appendage, an NTD and clathrin ankles. On binding, an auxilin DNAJ domain would overlap with the binding site for Eps15 on the β2-appendage, and would be able to interact with the NTD (consistent with (Schmid et al., 2006) by contacting the ‘Royle’ box binding groove (Willox and Royle, 2012) (**Fig. 4B,C**). Interactions between clathrin, auxilin, the β2-appendage and Eps15 at this regulatory nexus could be competitive or synergistic, coupling the concentration of adaptors (and therefore cargo) to dynamic rearrangement of the clathrin lattice. β2-appendage preferentially binds to hexagons, while bending of the clathrin lattice requires increasing the fraction of pentagons: hence expulsion of β2-appendage from its location in the coat and increasing lattice curvature will favour each other. Together these observations suggest that cargo binding, coat curvature, and dynamic assembly and disassembly of the coat, could be connected and controlled by reconfiguring low affinity, high avidity interactions between clathrin, adaptors and auxilin at a single site on the clathrin scaffold.

## Acknowledgements

This study made use of electron microscopes at EMBL, the MRC-LMB EM Facility and the Cryo-EM facility, University of Cambridge, Department of Biochemistry, as well as the high-performance computing resources at EMBL and LMB. We thank Wim Hagen, Dustin Morado, Dimitri Chirgadze and Steve Hardwick, and the staff of the facilities for support; Dustin Morado, William Wan and Aaron Tan for sharing and providing support with data processing scripts; Svetlana Dodonova for assistance in initial stages of sample optimisation and data processing; Frank Thommen, Andres Lindau, Michael Wahlers, Jake Grimmett and Toby Darling for supporting computational resources.

## Author contributions

OK, BTK, DJO and JAGB designed the project. OK performed in vitro budding reactions, cryo-ET, subtomogram averaging and model building with assistance from JAGB. VKD performed cryo-EM, single-particle reconstruction and model building. BTK and DJO designed constructs, purified proteins and performed biochemical validations. All authors interpreted data. OK, DJO and JAGB wrote the manuscript with assistance from BTK and VKD. DJO and JAGB obtained funding and managed the project.

## Materials and methods

### Preparation of reagents

To produce recombinant AP2 proteins, bacterial expression and purification, clathrin purification from porcine brain, liposome pelleting, lipopeptide coupling and liposome preparation were done as in Jackson et al., 2010 and Kelly et al., 2014. AP2 FLAP lacks the α subunit hinge and appendage (**Fig. S1**). Δβ2-appendage FLAP additionally lacks the β2-appendage, but is extended beyond the hinge region with an inert, unstructured region comprising 106 residues taken from the Enterobacteria phage f1 attachment protein G3P (residues 220-326, UniProtKB P69169), followed by a decahistidine tag. This extension facilitates purification of a fully intact β2 hinge region, which is otherwise cleaved.

For clathrin pulldown experiments, Glutathone S-transferase (GST)-tagged adaptors (for capture on glutathione sepharose beads) were constructed in pGEX-4T2 by genetically fusing fragments of β2 to GST. GST-β2hingeapp comprises the hinge and appendage regions of β2 fused to GST, as described previously (Kelly et al., 2014). GST-β2hinge comprises the extended β2 hinge as described above for the “Δβ2-appendage FLAP” construct, fused to GST. As a non-clathrin-binding control, the green fluorescent protein (GFP) was fused to GST. Fusion proteins were expressed and purified by standard techniques, using overnight expression at 22°C in BL21(DE3)pLysS cells.

All lipids were purchased from Avanti Polar lipids. The YxxΦ cargo signal was derived from TGN38 protein with sequence CKVTRRPKASDYQRL, and the [ED]xxxL[LI] signal was derived from phosphorylated CD4 with sequence CHRRRQAERM(SPhos)QIKRLLSEK. N-terminal cysteines were used for lipid coupling to maleimide lipids.

### Glutathione sepharose pulldown assays

Clathrin (0.8 µM) and adaptor (1.5 µM) were mixed in HKM buffer (25 mM Hepes pH 7.2, 125 mM potassium acetate, 5 mM magnesium acetate), total volume 100 µl, and incubated at 21°C for 30 minutes. 50 µl aliquots of glutathione sepharose beads (GE Healthcare) were washed four times in HKM/T buffer (HKM buffer supplemented with 0.05% Tween-20) before resuspension in the reaction mixtures. After a further 30 minutes of incubation at 21°C, with continuous gentle inversion to prevent settling, the beads were pelleted by centrifugation, supernatants were removed and retained for gel analysis, and the beads were washed four times in HKM/T before resuspension in 100 µl of HKM buffer. The pellet (’p’) and supernatant (‘s’) samples were supplemented with 33µl 4x SDS gel loading buffer, heated to 37°C for 30 minutes with occasional vortexing, and finally heated to 95°C for 5 minutes before analysis by SDS-PAGE (**Fig. S8B**).

### Single particle cryo-electron microscopy and model building

Purified AP2 core was reconstituted into physiological buffer HKM2 (10 mM Hepes KOH pH7.2, 120 mM KOAc, 2 mM MgCl_2_) with 0.5 mM DTT and applied to Quantifoil grids (R1.2/1.3, 300 Cu mesh). Grids were glow-discharged for 60 s at 25 mAmp using a Pelco EasiGlow before application of 3.5 µL of sample (0.4 mg/mL) and plunge-freezing using a Vitrobot Mark IV (FEI Company) operated at 4°C and 95% humidity. Data collection was carried out on a Titan Krios transmission electron microscope (FEI/Thermo Fisher) operated at 300 keV, equipped with a Falcon 3EC direct detector (FEI/Thermo Fisher) in counting mode. Automated data acquisition was performed using FEI/Thermo Fisher EPU software at a nominal magnification of 75,000, which corresponds to a pixel size of 1.065 Å per pixel. Dose-fractionated movies were acquired using 60 second exposures and 60 fractions at a dose of 0.54 e_-_/Å^2^/s with a defocus range of -1.9 to -3.1 µm (summarised in **Table S1**).

Data quality assessment, movie frame alignment, estimation of contrast-transfer function parameters, particle picking and extraction were carried out using Warp (Tegunov and Cramer, 2019). 120,000 particle images were extracted with a box size of 240 pixels and imported into CryoSPARC (Punjani et al., 2017) for 2D classification and *ab initio* model building. 3D refinement, filtering and sharpening (B factor -200) was carried out using the Homogeneous Refinement utility in CryoSPARC, including Fourier shell correlation (FSC) mask auto-tightening, followed by Local Resolution calculation by gold-standard FSC threshold (0.143 threshold) using the refined mask and an adaptive window factor of 25 (**Fig. S3A,B**).

The previously-determined ’closed’ conformation of the AP2 core complex (PDB ID 2vgl) was fitted into the final cryo-electron microscopy volume using UCSF Chimera (Pettersen et al., 2004) and the phenix.dock_in_map function of the Phenix Software suite (CC = 0.83) (Liebschner et al., 2019). The structure was refined using phenix.real_space_refine and found to have an overall RMSD = 1.147 versus the 2vgl coordinates with no major structural rearrangement (**Fig. S3B**, **Table S1**).

### Cryo-electron tomography sample preparation and data acquisition

To reconstitute AP2 membrane recruitment and Clathrin/AP2 membrane budding, purified AP2 FLAP and clathrin were incubated with synthetic liposomes. The liposomes contained 10% brain PtdIns(4,5)*P_2_* and 10% DOPS in a POPC/POPE (3:2) mixture, with 3% content of each cargo lipopeptide. *In vitro* budding reactions contained 1.4 μM of AP2, 0.7 μM clathrin and 0.5 mg/ml of 400 nm extruded liposomes in HKM2 buffer with 5 mM β-mercaptoethanol. AP2 and liposomes were incubated for 5 minutes at room temperature, clathrin was then added and the reaction was incubated for a further 15 minutes at room temperature followed by 10 minutes incubation at 37° C. For the reaction without clathrin, liposomes were prepared by extrusion through 200 nm filter, and the clathrin addition was omitted. 10 nm gold fiducial markers in HKM2 buffer was added to the reaction (1:10 fiducials to the reaction volume ratio), and 3 µl of this mixture was back side blotted for 3 seconds at relative humidity 98% and temperature 18° C on a glow-discharged holey carbon grid (CF-2/1-3C, Protochips), before plunge-freezing in liquid ethane (Leica EM GP2 automatic plunger). Dose-symmetrical tilt series acquisition (Hagen et al., 2017) was performed on an FEI Titan Krios electron microscope operated at 300 kV a Gatan Quantum energy filter with a slit width of 20eV and a K2 direct detector operated in counting mode. The total exposure of ∼130 e^-^/A^2^ was equally distributed between 41 tilts. 10 frame movies were acquired for each tilt. The details of data collection are given in **Table S2**.

### Raw image processing and tomogram reconstruction

The raw movies were corrected for detector gain and pixel defects, aligned and integrated using alignframes from the IMOD package (Kremer et al., 1996). A few tilt series with tracking errors or large beam-induced sample movements were discarded. In addition, a small number of defective high-tilt images (identified by blur, tracking error, large objects like grid bar or contaminations coming in the field of view) were discarded. Tilt-series were low pass filtered according to the cumulative radiation dose (Grant and Grigorieff, 2015) and aligned using fiducial markers in the IMOD package. Non-contrast-transfer-function (CTF) corrected, 4-times binned tomograms, were reconstructed by weighted back-projection in IMOD. For 3D CTF-corrected tomograms, per tilt defocus estimation was performed in CTFPLOTTER on non-dose-filtered tilt series, and correction and reconstruction were done using novaCTF (Turonova et al., 2017) with 15 nm strip width. Tomograms were binned by 2, 4 and 8 times (hereafter called bin2, bin4 and bin8 tomograms) with anti-aliasing.

### Subtomogram alignment

Subtomogram alignment and averaging was performed using MATLAB (MathWorks) functions adapted from the TOM (Forster et al., 2005), AV3 (Nickell et al., 2005), and Dynamo packages (Castano-Diez et al., 2012) essentially as described previously, using a modified wedge mask representing the amplitudes of the determined CTF and applied exposure filters at each tilt (Schur et al., 2016; Wan et al., 2017). **Table S3** summarises data processing parameters.

#### Tracing decorated membrane and initial subtomogram positions

To define initial positions of subtomograms, centres and radii of coated vesicles, buds and tubules were manually marked in bin4 tomograms using a Chimera plug-in (Qu et al., 2018). These measurements were used to define geometrical shapes: flexible tubes (for coated tubules) and spheres (for coated buds and vesicles). Subtomogram positions then were defined on the surface of these shapes (for AP2) or 120 Å above it (for clathrin) with uniform sampling at every ∼45 Å or ∼110 Å for AP2 and clathrin respectively. Initial subtomogram orientations were calculated to be normal to the shapes’ surfaces with random in-plane rotation.

#### Ab initio reference generation

Subtomograms were extracted at the initial positions from bin8 tomograms and averaged according to their initial orientations. Subtomograms were then aligned to this average in the direction perpendicular to the membrane and averaged to generate a reference in which density layers corresponding to the lipid bilayer, adaptor and clathrin layers were visible.

A subset of the data (three tomograms) was then aligned against this reference. For AP2, a gaussian-blurred sphere 80 Å in diameter was first added to the adaptor layer to assist the convergence of alignment and a cylindrical mask was applied passing the AP2 layer and the membrane. Three iterations of alignment were performed with a 10° angular search increment and a 35 Å low pass filter. For clathrin a cylindrical mask was applied passing the clathrin layer and excluding the membrane layer. Four iterations of alignment were performed with a 5° angular search increment and a 47 Å low pass filter. The resulting average was shifted, rotated and 3-fold symmetrised to place the clathrin triskelia in the centre of the box. In this manner separate starting references were generated for AP2 and clathrin.

#### Subtomogram alignment

These references were then used to align the complete data sets with the same alignment parameters. Upon alignment convergence, oversampling was removed by selecting a single subtomogram with the highest cross-correlation score within a distance threshold of 71 Å or 100 Å for AP2 and clathrin respectively.

Subtomograms were then sorted into identically sized odd and even subsets of coated structures. Subtomograms were extracted from bin4 tomograms and subsequent alignment was performed independently on the odd and even subsets. The search space and increments for angular and spatial parameters were gradually decreased, and the low pass-filter was gradually moved towards higher resolution. Upon convergence of subtomograms from bin4 tomograms, alignment was continued using bin2 and finally bin1 tomograms. Visibly misaligned subtomograms were removed using a cross-correlation cut-off threshold manually selected for each tomogram. Resolution was monitored by FSC. The final maps were sharpened with empirically determined B factor with local low-pass filtering using relion_postprocess from the Relion 3.0 package (Zivanov et al., 2018).

For AP2 alone, after convergence, focussed alignment was performed by local masking of either α, β2/µ2 or the C-termini of α and β2. The focussed maps were multiplied by their alignment mask, summed, and divided by the sum of all three local masks to obtain the final EM map. Focussed alignment was not performed for AP2 in clathrin coats due to insufficient signal. Final maps were multiplied by a soft cylindrical mask to remove diffuse density from neighbouring AP2 molecules.

For clathrin, prior to alignment in bin2 tomograms, we performed symmetry expansion using dynamo_table_subbox from the Dynamo package (Castano-Diez et al., 2012) to generate subtomograms centered at individual clathrin legs and further alignment iterations were performed on individual legs.

#### Classification of subtomograms

We applied principal component analysis (PCA) to sort subtomograms by β2-appendage occupancy. The PCA was performed on wedge-masked difference maps (Heumann et al., 2011) with calculations implemented in MATLAB using code adapted from PEET and Dynamo packages (Castano-Diez et al., 2012; Heumann et al., 2011). The first eigencomponent of this PCA analysis correlated with the β2-appendage density in the subtomograms and could be used to sort the data according to β2-appendage occupancy. We sorted subtomograms bounding pentagons or hexagons of the clathrin lattice based on the positions of their neighbours (**Fig. S5A**).

To produce a β2-appendage-enriched subtomogram class, hexagon- and pentagon-bounding subtomograms were split into 10 equally-sized classes according to β2-appendage occupancy (**Fig. S6A**). The four classes of hexagon-facing legs with highest β2-appendage occupancy were combined together and the combined class was used for further alignment to derive the EM map resolving the β2-appendage density in the clathrin cage.

All EM maps generated from the two clathrin datasets (with YxxΦ and YxxΦ/[ED]xxxL[LI] cargoes) were essentially identical. Both clathrin datasets were therefore combined for further alignment in bin2 and bin1.

### Spatial distributions of AP2 and Clathrin

Plots showing the distribution of proteins relative to their neighbors were created as in (Kovtun et al., 2018). Plots of the distribution of AP2 relative to AP2 (**Fig S6**), relate the centers of AP2 complexes. These plots were used to identify AP2 dimers in the dataset of AP2 on YxxΦ cargo-containing membrane which appear as a peak in the distribution containing ∼20 % of total AP2 (marked by arrow in **Fig. S4A**). To derive a structure of the dimer, subtomograms containing dimers were aligned in bin4 without symmetrisation then in bin2 with 2-fold symmetrisation until convergence.

Plots of the distribution of clathrin relative to clathrin (**Fig. S6**), relate the CHCR3 regions of distal clathrin segments. These plots were used to separate clathrin legs that bound hexagons and pentagons in the clathrin cage (**Fig. S6**).

Plots of the distribution of AP2 relative to clathrin, relate the center of AP2 and the tripod region of the clathrin leg. Plots of the distribution of clathrin relative to AP2 relate the center of AP2 and the membrane-facing edge of clathrin NTDs. These plots were used to analyse spatial relationships between the adaptor and clathrin components of the coat (**Fig. S6**).

Linear profiles of the radial distribution of AP2 relative to the clathrin tripod were derived by integrating the AP2 distribution plot along the axis of a cylindrical mask with a radius of 54 Å. The mean and standard deviation of the linear profiles were calculated by gaussian fitting (Python, Matplotlib).

### Model building in EM maps determined by subtomogram averaging

Rigid-body docking was performed using the Chimera package (Pettersen et al., 2004). Flexible fitting was performed using molecular dynamics flexible fitting (MDFF) in the NAMD package (Trabuco et al., 2008), constraining secondary structure elements. For AP2, fitting was performed using the crystal structure of the open form (PDB ID 2xa7)(Jackson et al., 2010). For clathrin light chain fitting was performed using (PDB ID 6sct) (Morris et al., 2019). For clathrin heavy chain, fitting was performed using (PDB ID 1xa4) (Fotin et al., 2004b). PDB ID 1xa4 is a C-alpha trace model based upon EMDB entry EMD-5119, and to allow MDFF, side chains were added to the model using PD2 ca2main online server (Moore et al., 2013). The separate fragments in PDB ID 1xa4 (1-834, 840-1135, 1078-1278, 1281-1630) were fitted as rigid bodies into the map of a clathrin leg bounding a hexagon, joined using Modeller (Sali and Blundell, 1993), and combined with the rigid-body fitted light chain to make a composite model of CHC and CLC, that was further refined by flexible fitting with MDFF. The resulting refined model was subsequently flexibly fitted into a map of leg bounding a hexagon enriched in the β2-appendage, constraining NTDs as a single domain in addition to the secondary structure constraints used above. This was done to refine the positions of the clathrin NTD-ankle regions, that are better resolved in the β2-appendage-enriched map compared to the map representing all hexagon bounding legs. The β2-appendage (PDB ID 1e42, (Owen et al., 2000) was fit as a rigid body using Chimera. To display the position of the DNAJ domain of Auxilin, the model of NTD/DNAJ taken from PDB ID 1xi5 (Fotin et al., 2004a) was overlaid with the clathrin NTD from our pseudoatomic model.

---

**Fig. S1.**
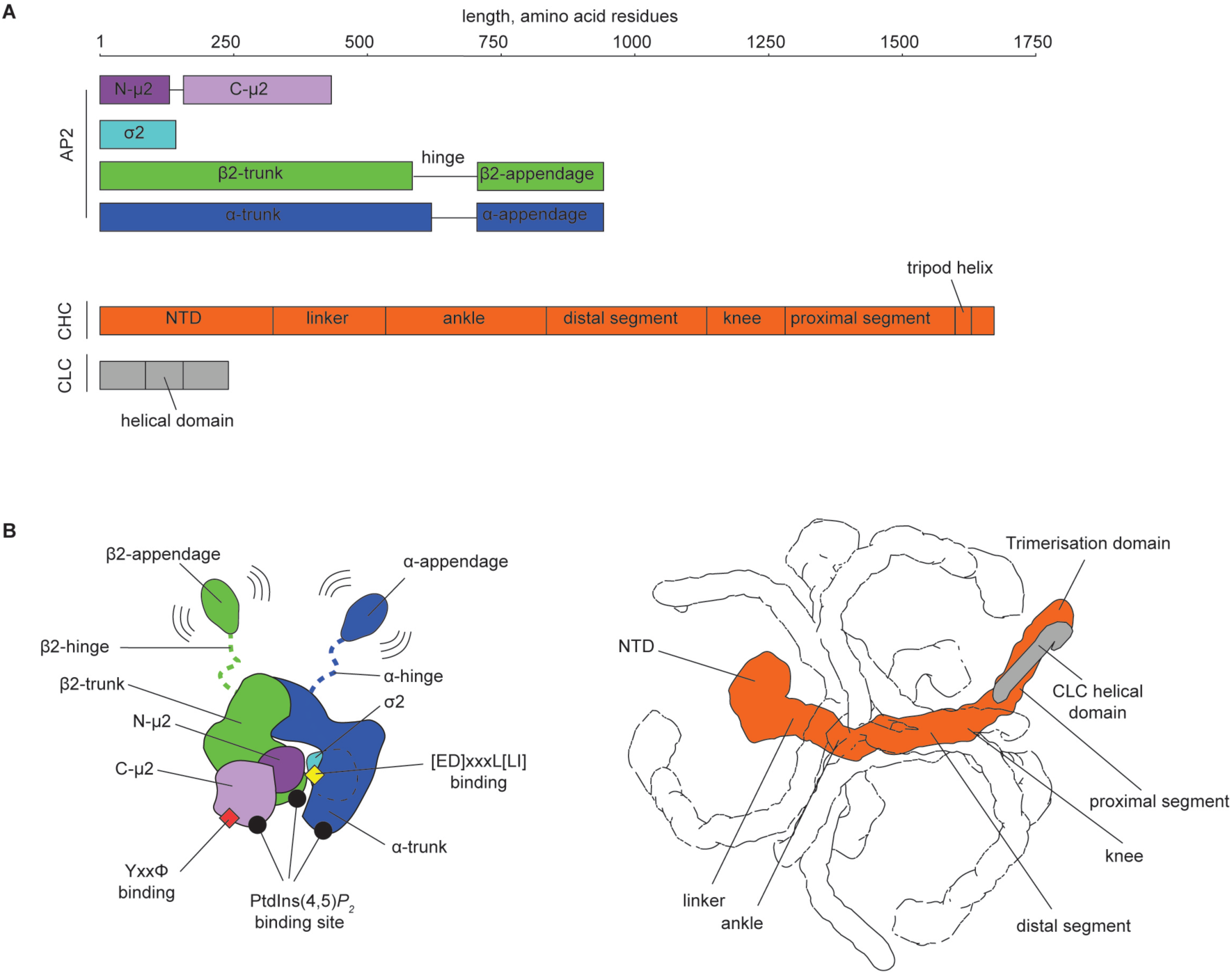
Overview of AP2 and clathrin structures. **A.** Schematic representation of AP2 and clathrin polypeptide chains with marked domains. A length ruler (in amino acid residues) is given at the top. Note that the “FLAP” AP2 construct lacks α hinge and appendage regions, while the “Core” AP2 construct lacks α and β2 hinge and appendage regions. **B**. Cartoon representation of the structures of AP2 (left) and Clathrin (right). Subunits are color-coded as in **A** and known functional sites and domains are marked. For clathrin, neighboring heavy chain subunits within a triskelia are shown as outlines.

**Fig. S2.**
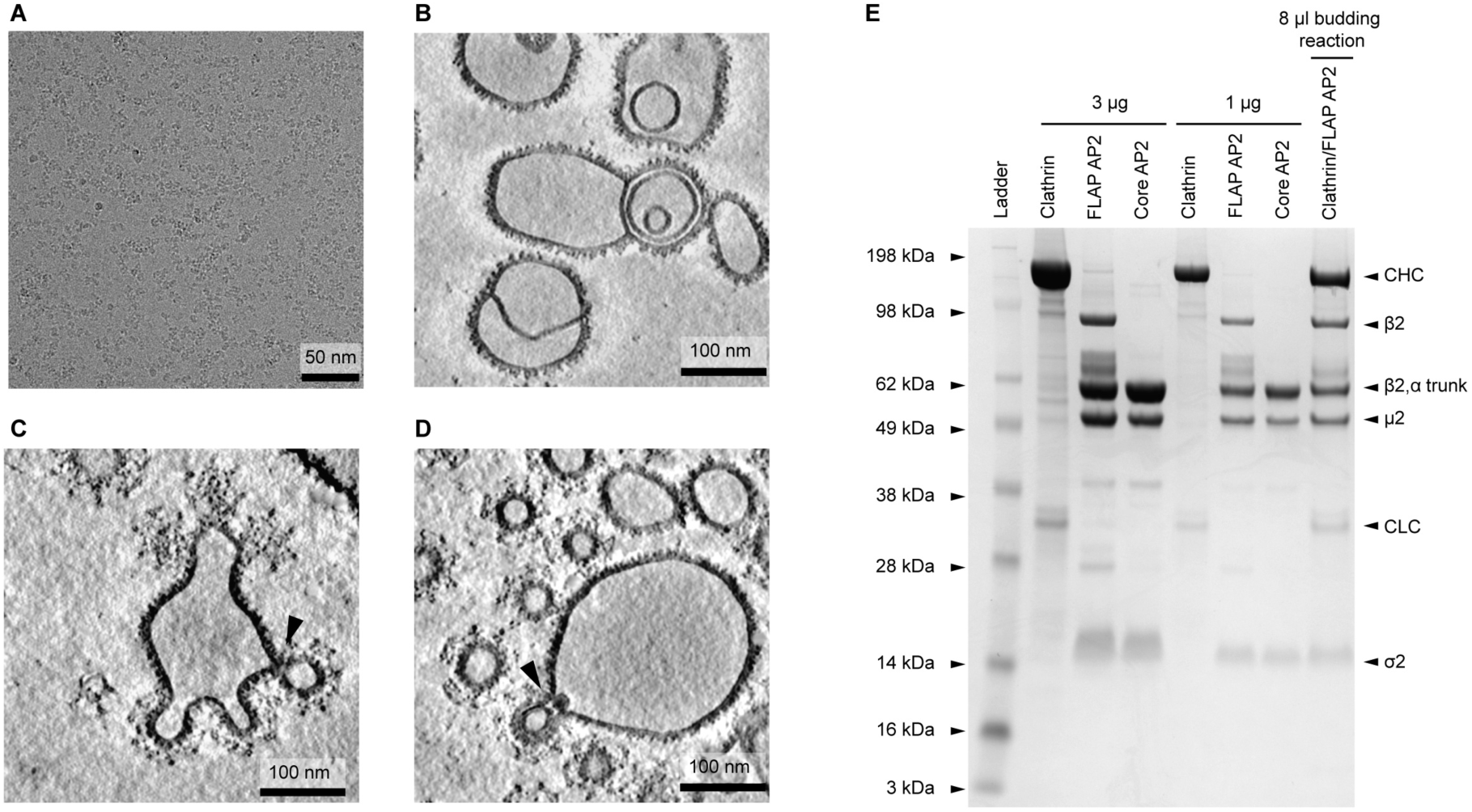
Representative EM images of datasets used for structural characterisation and PAGE characterisation of protein preparations and budding reactions. **A.** A cryo-EM image of AP2 in buffer mimicking physiological conditions (see methods). **B.** Slices through a tomogram of AP2 recruited to membranes containing YxxΦ cargo. **C.** Clathrin/AP2 coats assembled on membranes containing YxxΦ cargo. **D.** Clathrin/AP2 coats assembled on membranes containing YxxΦ and [ED]xxxL[LI] cargo. In **B-D**, AP2 appears as a dense speckled coat on the membrane. Clathrin forms polygonal cages around the highly curved membranes of buds and vesicles. Coated buds often have narrow necks (arrowheads). Small vesicles only appear upon the addition of clathrin, suggesting that clathrin-coated vesicles formed via scission of their narrow necks. **E.** Electrophoretic separation in 4-12% NuPAGE gel in MOPS buffer (Invitrogen) of purified clathrin, AP2 and the complete budding reaction along with a molecular size ladder. FLAP AP2 lacks α hinge and appendage regions. Core AP2 lacks α and β2 hinge and appendage regions.

**Fig. S3.**
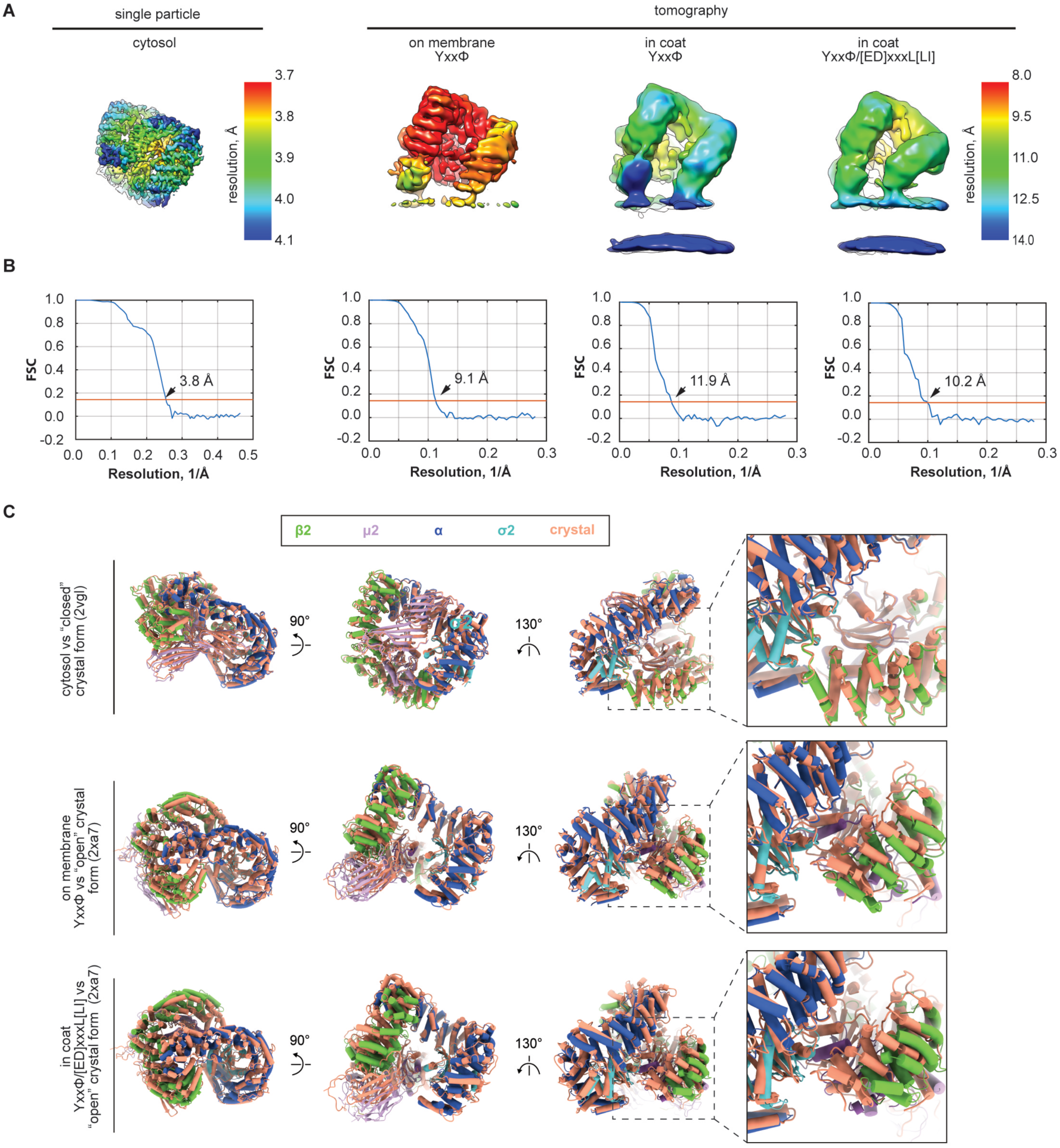
Resolution assessment of AP2 EM maps. Overlay of ribbon models of AP2 in physiological conditions and related crystal structures. **A.** EM maps colored by the local resolution for the four AP2 forms determined in this work. **B.** The corresponding FSC plots with arrows indicating the resolution at 0.143 FSC criterion. **C**. Overlays of AP2 structures modelled based on the EM maps in **A** and related crystal structures of the closed (2vgl, (Collins et al., 2002) and open (2xa7,(Jackson et al., 2010) forms. Panels in the right column show close up views of the N-terminal region of β2. The β2 position is essentially the same in the cytosolic and closed crystal form of AP2, but appears more open in membrane-bound AP2 than in the “open” crystal form.

**Fig. S4.**
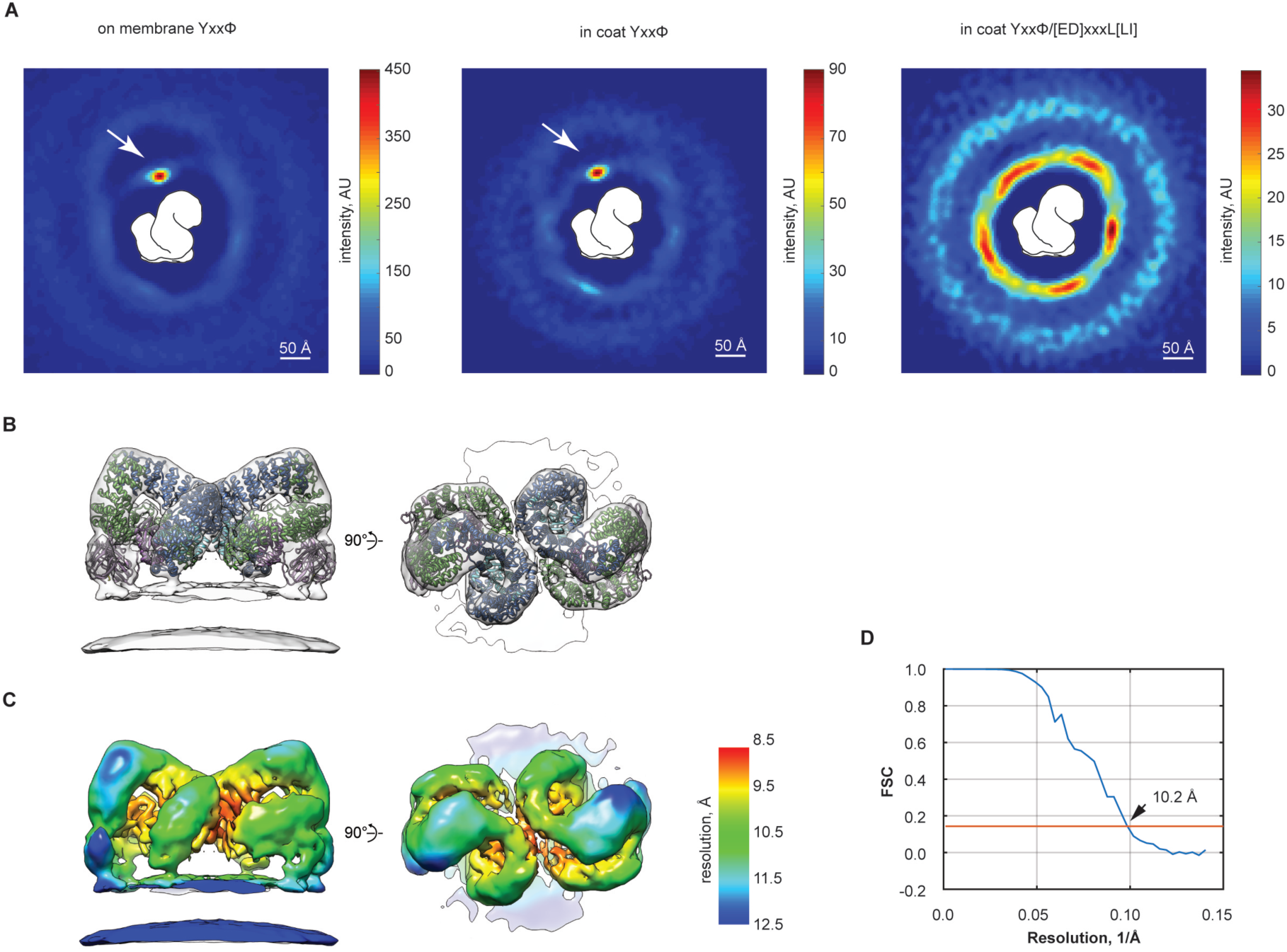
Distribution of neighboring AP2s on membranes. **A.** Plots (“neighbor plots”) show the spatial distribution of the centers of neighbouring AP2 molecules for the entire dataset of a specific type. High intensity indicates that a neighbouring AP2 molecule is found with high frequency at that position in space relative to the central AP2 molecule (outlined in white). The neighbor plot is three dimensional - for visualisation, a 60 Å thick central slab of the neighbor plots has been integrated perpendicular to the membrane and coloured by intensity. Arrows indicate preferred positions of neighbors corresponding to an AP2 dimer. The dimer is frequent on membranes containing YxxΦ-cargo (**left** and **center panels**) but not membranes containing YxxΦ/[ED]xxxL[LI] cargo (**right panel**). **B**. Averaging all AP2 molecules which have a neighbor at the dimer position generates a structure of the dimer. The EM structure of the dimer (transparent isosurface) has been rigid-body-fitted with two copies of the structural model for AP2 on YxxΦ-cargo membranes. **C.** The EM map of the AP2 dimer colored by local resolution. **D.** the corresponding FSC plot.

**Fig. S5.**
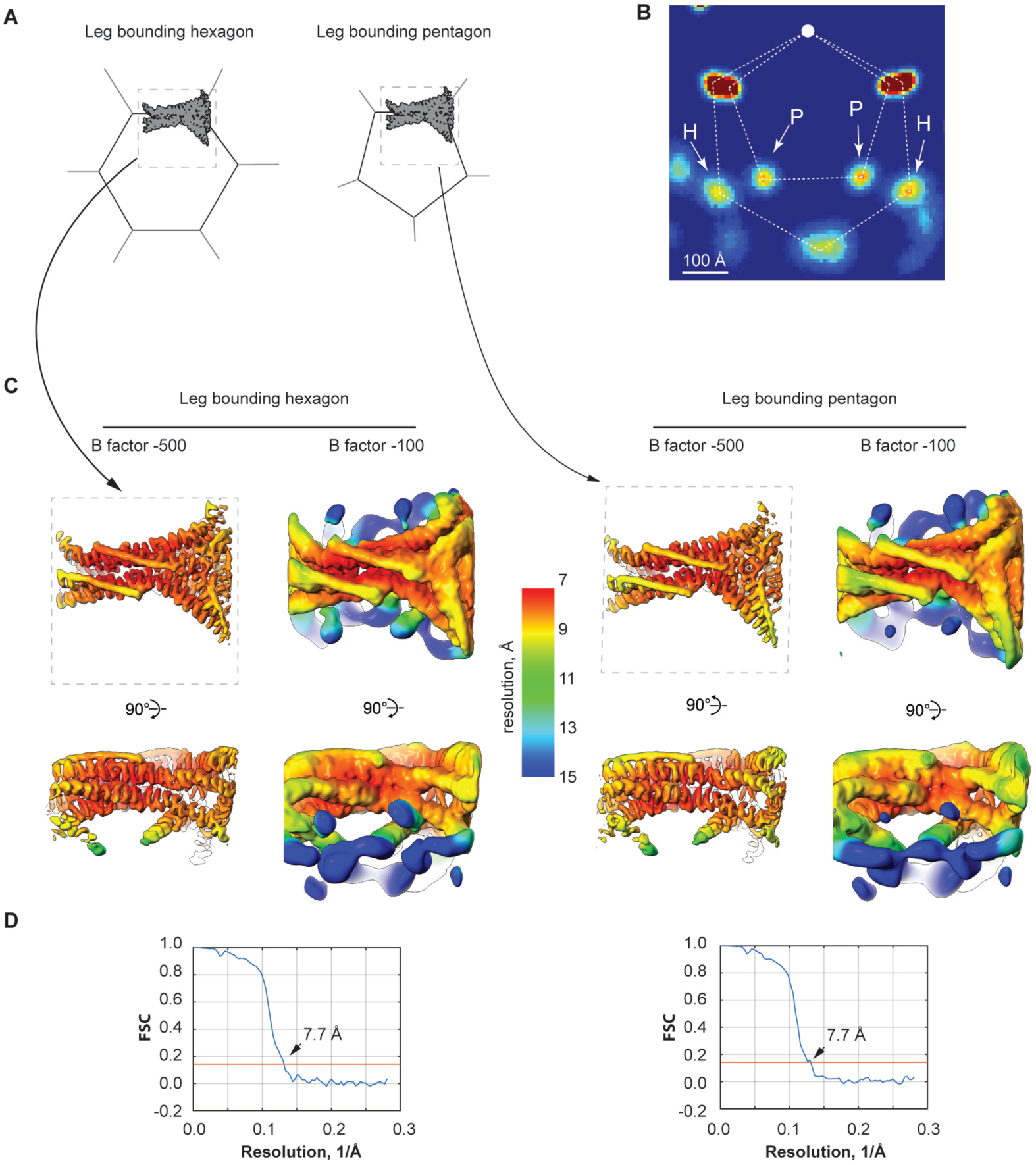
EM maps of clathrin legs bounding hexagons and pentagons. **A.** A schematic representation of subtomograms containing a clathrin leg bounding a hexagon (**left**) or a pentagon (**right**) of the clathrin cage. The dotted boxes indicate boundaries of subtomograms with a cartoon representation of the clathrin EM map from **C** (indicated by long arrows). **B.** Neighbor plot showing the spatial distribution of neighboring clathrin legs in the YxxΦ /[ED]xxxL[LI] coat (visualised as in **S4A** but integrating over a ∼110 Å-thick slab). The white disc indicates the origin relative to which positions of neighbors were determined. Dotted lines outline clathrin polygons. Arrows indicate peaks where neighboring legs are frequently present marked according to whether they correspond to neighbors around a hexagon (H) or a pentagon (P). Clathrin legs were sorted into hexagon and pentagon bounding classes according to whether their neighbours were at H or P positions. **C.** EM maps of the hexagon- and pentagon-bounding classes combined from both YxxΦ and YxxΦ /[ED]xxxL[LI] coat datasets, coloured by resolution as in **Fig. S3A**. The maps are shown at different levels of sharpening to reveal secondary structure (B factor -500) or low-resolution features such as NTD domains and β2-appendages (B factor -100). The resolution drops sharply in the NTD-ankle region, demonstrating the mobility of these clathrin elements compared to the rest of the structure. **D.** The corresponding FSC plots.

**Fig. S6.**
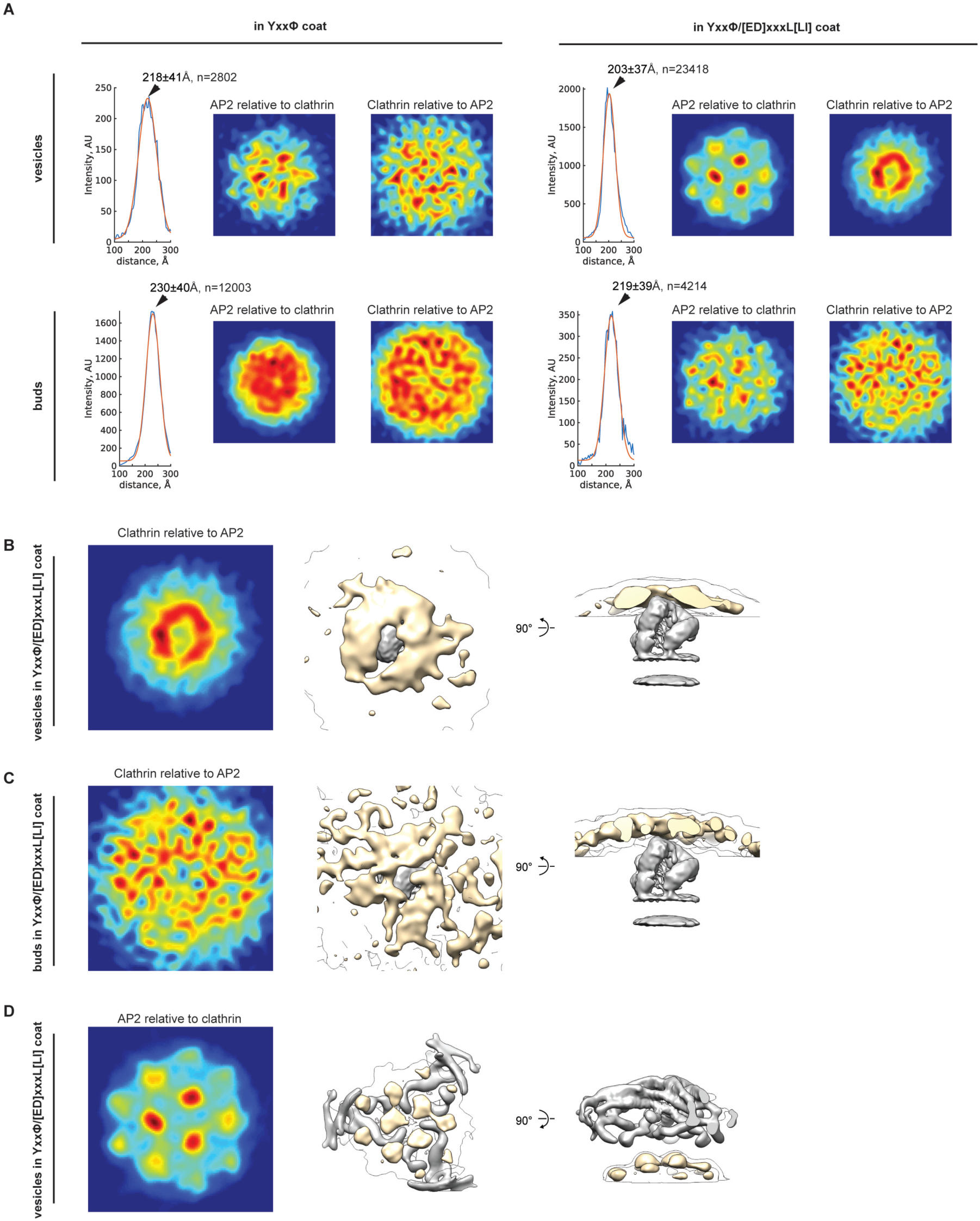
Spatial relationship between AP2 and clathrin in coats. **A**. Neighbor plots showing the distribution of the center of AP2 relative to the clathrin tripod or the distribution of clathrin NTDs relative to the center of AP2 molecules. Plots are visualized as in **Fig S4A**, integrating 90 Å-thick slabs (for AP2 relative to clathrin) or 82 Å-thick slabs (for clathrin relative to AP2). Also shown are distributions of the radial distance between AP2 and clathrin calculated from the “AP2 relative to clathrin” neighbor plots (blue lines, see supplementary methods). Mean +/-standard deviation determined by fitting with a Gaussian distribution (red lines) and the number of AP2 subtomograms considered is marked. The distance between clathrin and AP2 is shorter in YxxΦ/[ED]xxxL[LI] cargo-containing coats than in YxxΦ cargo-containing coats, and is shorter in vesicles than in buds. AP2 begins to occupy preferred positions between clathrin NTDs at a distance between AP2 and clathrin shorter than ∼220 Å, and this preference becomes pronounced at a distance of ∼203 Å as seen in YxxΦ /[ED]xxxL[LI] coated vesicles. Corresponding changes are seen in the distribution of clathrin relative to AP2, with clathrin NTD being excluded from positions directly above AP2 molecules (see also panel **B**). **B-D.** Selected neighbor plots from **A** are additionally illustrated as isosurfaces (the beige isosurface encloses 8% of relative subtomogram positions, the transparent outline encloses 40%), relative to the protein structures (grey surface). **B.** Clathrin NTDs are preferentially excluded from regions directly above AP2 in vesicles where the distance between AP2 and clathrin is short enough for them to sterically exclude one another. **C.** Clathrin NTDs are randomly distributed in buds where the distance between AP2 and clathrin is large enough to avoid any steric exclusion. **D.** The preferred positions of AP2 coincide with the spaces between the clathrin NTDs – each space provides enough room to accommodate the tip of a single AP2.

**Fig. S7.**
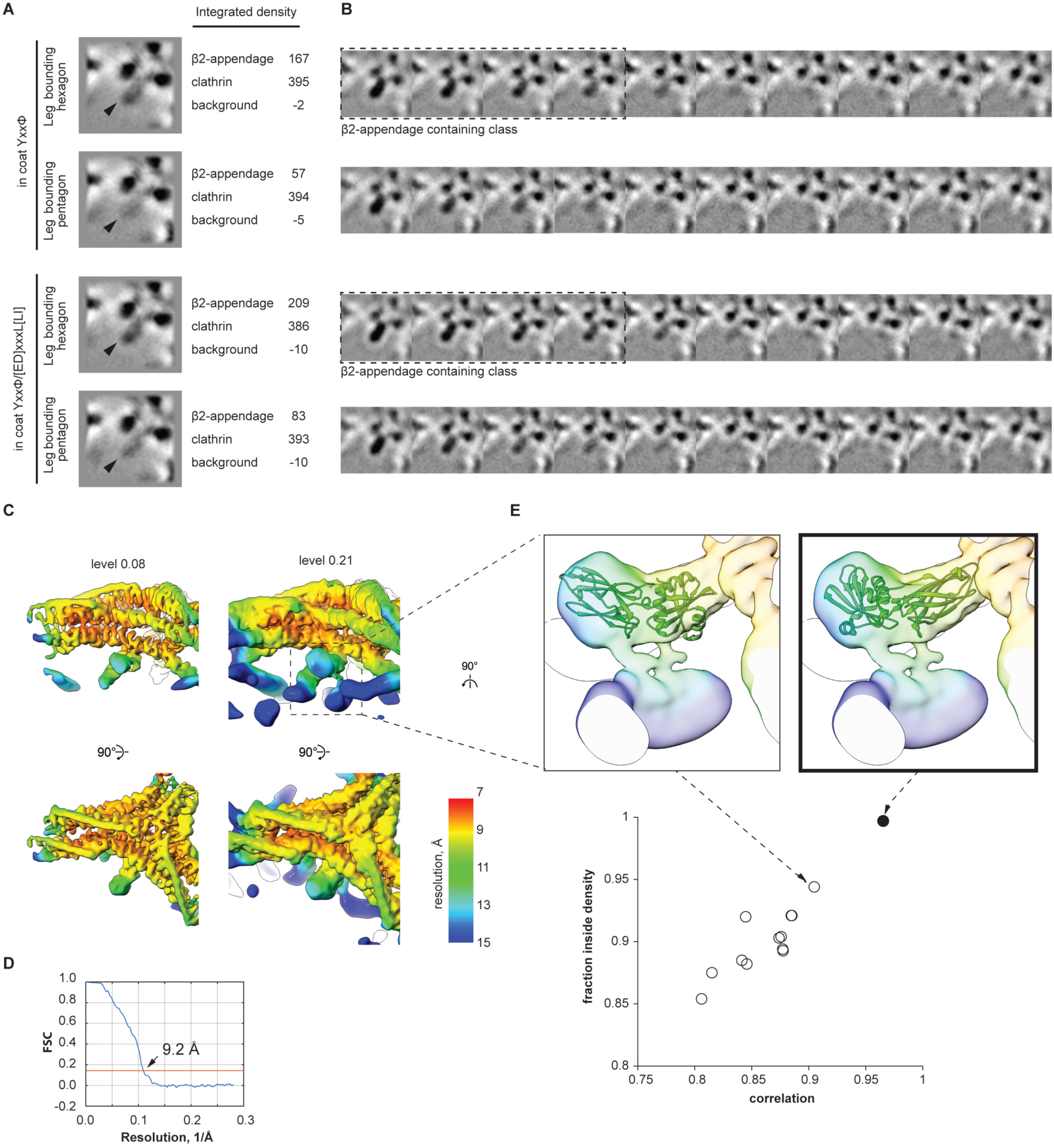
Classification of β2-appendage enriched subtomograms and rigid body fitting of the atomic structure of β2-appendage. **A**. Slices through EM maps of clathrin legs bounding hexagons and pentagons. The β2-appendage density (arrowheads) is stronger in the hexagon-bounding leg than in the pentagon-bounding leg. The pixel density was integrated within an ellipsoidal mask placed on the β2-appendage, clathrin, or background, revealing higher occupancy if the β2-appendage on legs bounding hexagons than bounding pentagons. We estimate that the occupancy is approximately 50% and 20% in legs bounding hexagons and pentagons, respectively. **B**. Subtomograms classified based on β2-appendage occupancy. Panels are arranged in rows by the polygon and coat types as in **A**. The datasets were split into ten equally-sized classes by the score of the principal component that represents the β2-appendage density. Classes of hexagon-bounding legs enriched by β2-appendage were selected using visual inspection (classes within dotted boxes), and the corresponding subtomograms were combined into a single dataset which was averaged to produce the EM map shown in **C**. **C**. EM map of clathrin leg bounding hexagon enriched in the β2-appendage. The map is shown at two different isosurface levels and is coloured by local resolution as in **Fig. S3A**. **D.** Corresponding FSC plot. **E.** Rigid body fit of the atomic model of the β2-appendage domain (PDB ID 1e42, (Owen et al., 2000) into the EM map from **C** was assessed based on correlation and the fraction of the model within the density, revealing an outlying highest-scoring fit (filled circle, **lower panel**). The EM map and the fitted ribbon models for the highest-scoring and second highest-scoring fits are illustrated. Only the highest scoring fit is consistent with the EM density (bold frame).

**Fig. S8.**
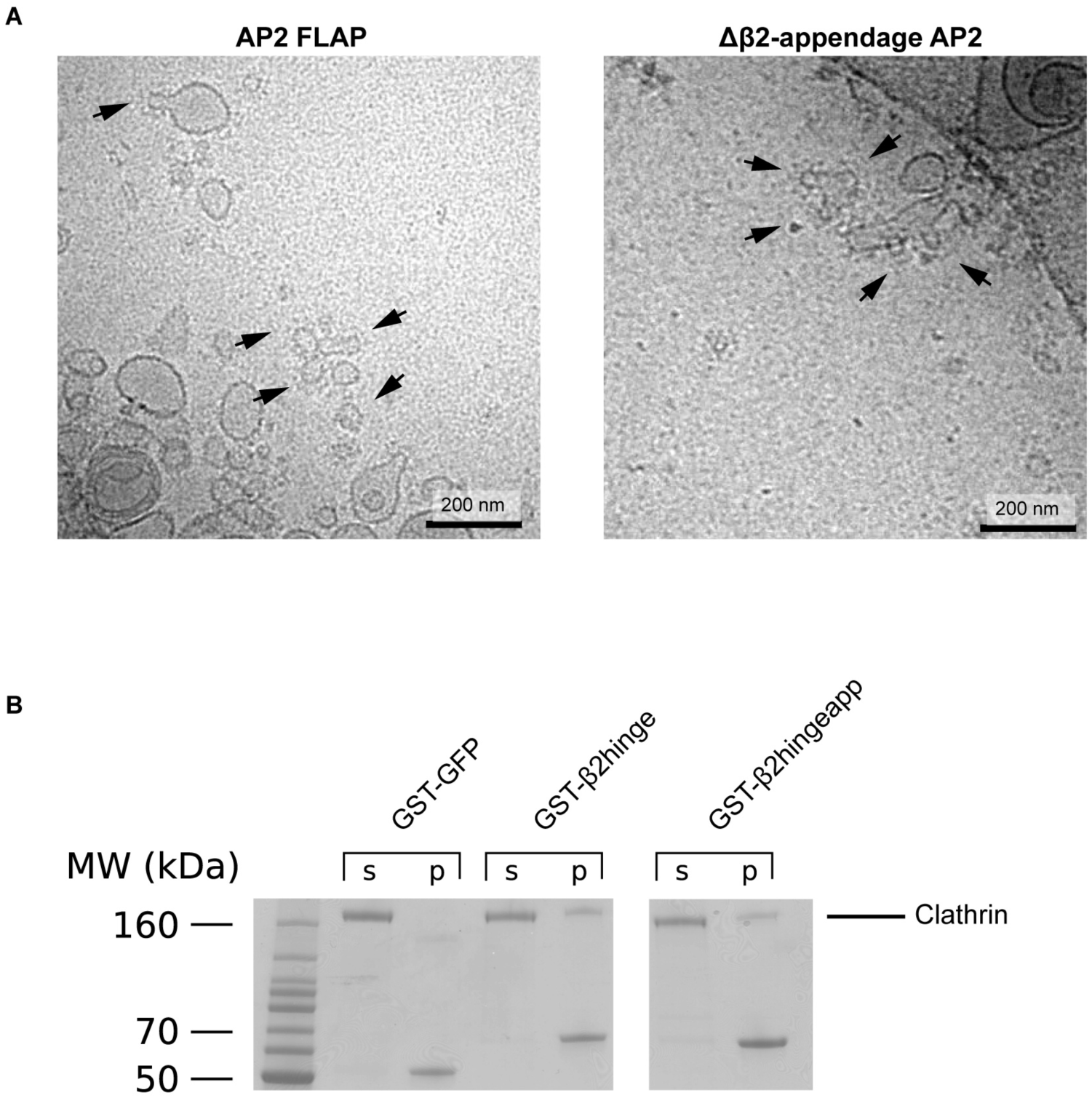
Deletion of β2-appendage in AP2 FLAP does not preclude the formation of clathrin-coated buds. **A.** Cryo-EM screening images showing budding reactions on YxxΦ cargo-containing liposomes using AP2 FLAP and AP2 FLAP lacking the β2-appendage (Δβ2-appendage). Arrows indicate clathrin-coated buds formed in the budding reactions. **B.** Coomassie stained SDS-PAGE of glutathione sepharose pulldowns using GST-GFP (as a negative control for clathrin binding), GST-β2hinge or GST-β2hingeapp. Supernatant (‘s’) and pellet (‘p’). GST-β2hinge and GST-β2hingeapp bind clathrin to a similar extent, suggesting that deletion of the β2-appendage does not affect clathrin recruitment in this system.

**Table S1.**
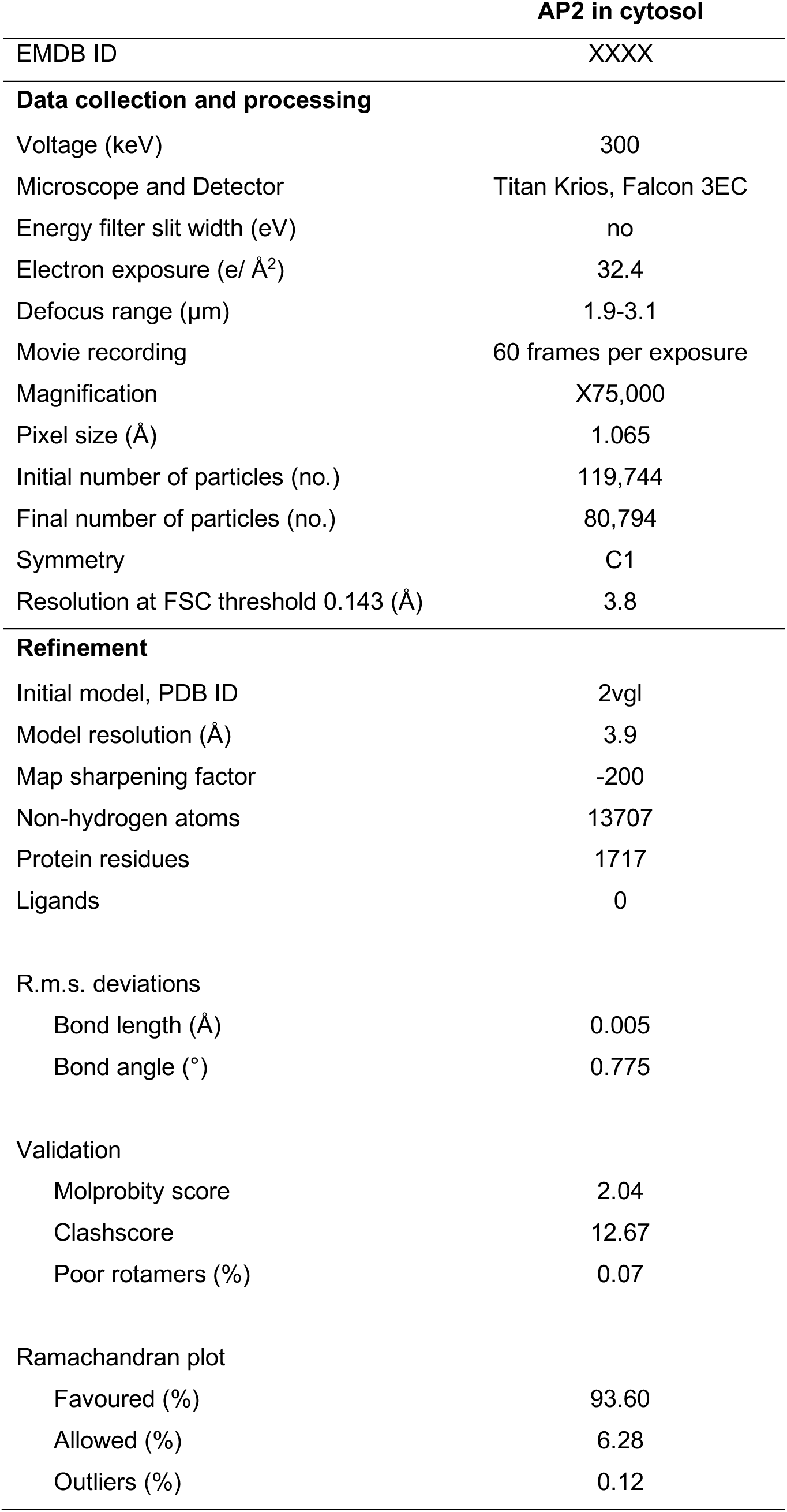
Cryo-electron microscopy data collection, refinement and validation statistics for single-particle structure of AP2 in solution.

**Table S2.** Cryo-electron tomography data collection parameters for membrane-recruited AP2 and clathrin.

|  | AP2<br>YxxΦ membrane | AP2/Clathrin<br>YxxΦ membrane | AP2/Clathrin<br>YxxΦ/[ED]xxxL[LI]<br>membrane |
| --- | --- | --- | --- |
| Voltage (keV) |  | 300 |  |
| Microscope and Detector |  | Titan Krios, Gatan Quantum K2 |  |
| Energy filter slit width (eV) |  | 20 |  |
| Electron exposure (e/Å<sup>2</sup>) dose fractionation |  | ~130, uniformly distributed over tilt series |  |
| Defocus range (μm) | 2.5-5.5 | 2.5-6.5 | 1.5-4.5 |
| Tilt scheme (min/max, step) |  | -60°/+60, 3°, dose-symmetrical (Hagen scheme) |  |
| Movie recording |  | 10 frames per tilt |  |
| Magnification (times) |  | X81,000 |  |
| Pixel size (Å) | 1.780 | 1.780 | 1.787 |
| Number of tomograms acquired/used (no.) | 77/66 | 48/45 | 44/43 |
| Traced membranes (no.) and lumenal diameter<br>(mean±std) | 1083 liposomes (99±46 nm)<br>244 tubules (19±7 nm) | 397 buds (35±4 nm)<br>55 vesicles (70±10 nm) | 96 buds (56±6 nm)<br>564 vesicles (56±10nm) |

**Table S3.** Subtomogram averaging image processing parameters and statistics for membrane-recruited AP2 and clathrin.

|  | AP2 |  |  |  | Clathrin |  |  |
| --- | --- | --- | --- | --- | --- | --- | --- |
|  | on YxxΦ membrane |  | in YxxΦ coat | in YxxΦ/<br>[ED]xxxL[LI] | hexagon-bounding | pentagon-bounding | hexagon-bounding, β2-appendage<br>enriched |
|  | monomer | dimer |  | coat |  |  |  |
| EMDB ID | XXXX | XXXX | XXXX | XXXX | XXXX | XXXX | XXXX |
| Initial subtomograms (no.) | 585,669 |  | 224,577 | 292,224 |  | 79,913<sup>1</sup>/118,958<sup>2</sup> |  |
| Subtomograms after distance/cluster<br>cleaning (no.) | 112,237 |  | 25,694 | 29,139 |  | 20,917<sup>1</sup>/37,204<sup>2</sup> |  |
| Subtomograms after cross-correlation<br>threshold cleaning (no.) | 100,556 |  | 14,805 | 27,630 |  | 18,371<sup>1</sup>/23,832<sup>2</sup> |  |
| Subtomograms after symmetry<br>expansion (no.) | n/a |  | n/a | n/a |  | 55,131<sup>1</sup>/71,496<sup>2</sup> |  |
| Final subtomogram (no.) | 100,556 | 22,323 | 14,805 | 27,630 | 30,190 | 24,759 | 12,076 |
| Symmetry | C1 | C2 | C1 | C1 | C1 | C1 | C1 |
| Resolution at FSC threshold 0.143 (Å) | 9.1 | 10.2 | 11.9 | 10.2 | 7.7 | 7.7 | 9.2 |
| Resolution range (Å) | 7-18 | 8.5-24 | 9-24 | 9-24 | 6-25 | 6-25 | 7.5-30 |

1. YxxΦ coat 2. YxxΦ/[ED]xxxL[LI] coat

## References

1. Borner, G.H., R. Antrobus, J. Hirst, G.S. Bhumbra, P. Kozik, L.P. Jackson, D.A. Sahlender, and M.S. Robinson. 2012. Multivariate proteomic profiling identifies novel accessory proteins of coated vesicles. J Cell Biol. 197:141–160.

2. Castano-Diez, D., M. Kudryashev, M. Arheit, and H. Stahlberg. 2012. Dynamo: a flexible, user-friendly development tool for subtomogram averaging of cryo-EM data in high-performance computing environments. J Struct Biol. 178:139–151.

3. Chen, Y., J. Yong, A. Martinez-Sanchez, Y. Yang, Y. Wu, P. De Camilli, R. Fernandez-Busnadiego, and M. Wu. 2019. Dynamic instability of clathrin assembly provides proofreading control for endocytosis. J Cell Biol. 218:3200–3211.

4. Cocucci, E., F. Aguet, S. Boulant, and T. Kirchhausen. 2012. The first five seconds in the life of a clathrin-coated pit. Cell. 150:495–507.

5. Collins, B.M., A.J. McCoy, H.M. Kent, P.R. Evans, and D.J. Owen. 2002. Molecular architecture and functional model of the endocytic AP2 complex. Cell. 109:523–535.

6. Dodonova, S.O., P. Diestelkoetter-Bachert, A. von Appen, W.J. Hagen, R. Beck, M. Beck, F. Wieland, and J.A. Briggs. 2015. VESICULAR TRANSPORT. A structure of the COPI coat and the role of coat proteins in membrane vesicle assembly. Science. 349:195–198.

7. Edeling, M.A., S.K. Mishra, P.A. Keyel, A.L. Steinhauser, B.M. Collins, R. Roth, J.E. Heuser, D.J. Owen, and L.M. Traub. 2006. Molecular switches involving the AP-2 beta2 appendage regulate endocytic cargo selection and clathrin coat assembly. Dev Cell. 10:329–342.

8. Forster, F., O. Medalia, N. Zauberman, W. Baumeister, and D. Fass. 2005. Retrovirus envelope protein complex structure in situ studied by cryo-electron tomography. Proc Natl Acad Sci U S A. 102:4729–4734.

9. Fotin, A., Y. Cheng, N. Grigorieff, T. Walz, S.C. Harrison, and T. Kirchhausen. 2004a. Structure of an auxilin-bound clathrin coat and its implications for the mechanism of uncoating. Nature. 432:649–653.

10. Fotin, A., Y. Cheng, P. Sliz, N. Grigorieff, S.C. Harrison, T. Kirchhausen, and T. Walz. 2004b. Molecular model for a complete clathrin lattice from electron cryomicroscopy. Nature. 432:573–579.

11. Gaidarov, I., Q. Chen, J.R. Falck, K.K. Reddy, and J.H. Keen. 1996. A functional phosphatidylinositol 3,4,5-trisphosphate/phosphoinositide binding domain in the clathrin adaptor AP-2 alpha subunit. Implications for the endocytic pathway. J Biol Chem. 271:20922–20929.

12. Grant, T., and N. Grigorieff. 2015. Measuring the optimal exposure for single particle cryo-EM using a 2.6 A reconstruction of rotavirus VP6. Elife. 4:e06980.

13. Hagen, W.J.H., W. Wan, and J.A.G. Briggs. 2017. Implementation of a cryo-electron tomography tilt-scheme optimized for high resolution subtomogram averaging. J Struct Biol. 197:191–198.

14. Helbig, I., T. Lopez-Hernandez, O. Shor, P. Galer, S. Ganesan, M. Pendziwiat, A. Rademacher, C.A. Ellis, N. Humpfer, N. Schwarz, S. Seiffert, J. Peeden, J. Shen, K. Sterbova, T.B. Hammer, R.S. Moller, D.N. Shinde, S. Tang, L. Smith, A. Poduri, R. Krause, F. Benninger, K.L. Helbig, V. Haucke, Y.G. Weber, E.-R.E.S.C. Euro, and G. Consortium. 2019. A Recurrent Missense Variant in AP2M1 Impairs Clathrin-Mediated Endocytosis and Causes Developmental and Epileptic Encephalopathy. Am J Hum Genet.

15. Heumann, J.M., A. Hoenger, and D.N. Mastronarde. 2011. Clustering and variance maps for cryo-electron tomography using wedge-masked differences. J Struct Biol. 175:288–299.

16. Hollopeter, G., J.J. Lange, Y. Zhang, T.N. Vu, M. Gu, M. Ailion, E.J. Lambie, B.D. Slaughter, J.R. Unruh, L. Florens, and E.M. Jorgensen. 2014. The membrane-associated proteins FCHo and SGIP are allosteric activators of the AP2 clathrin adaptor complex. Elife. 3.

17. Honing, S., D. Ricotta, M. Krauss, K. Spate, B. Spolaore, A. Motley, M. Robinson, C. Robinson, V. Haucke, and D.J. Owen. 2005. Phosphatidylinositol-(4,5)-bisphosphate regulates sorting signal recognition by the clathrin-associated adaptor complex AP2. Mol Cell. 18:519–531.

18. Jackson, L.P., B.T. Kelly, A.J. McCoy, T. Gaffry, L.C. James, B.M. Collins, S. Honing, P.R. Evans, and D.J. Owen. 2010. A large-scale conformational change couples membrane recruitment to cargo binding in the AP2 clathrin adaptor complex. Cell. 141:1220–1229.

19. Jia, X., E. Weber, A. Tokarev, M. Lewinski, M. Rizk, M. Suarez, J. Guatelli, and Y. Xiong. 2014. Structural basis of HIV-1 Vpu-mediated BST2 antagonism via hijacking of the clathrin adaptor protein complex 1. Elife. 3:e02362.

20. Kadlecova, Z., S.J. Spielman, D. Loerke, A. Mohanakrishnan, D.K. Reed, and S.L. Schmid. 2017. Regulation of clathrin-mediated endocytosis by hierarchical allosteric activation of AP2. J Cell Biol. 216:167–179.

21. Kelly, B.T., S.C. Graham, N. Liska, P.N. Dannhauser, S. Honing, E.J. Ungewickell, and D.J. Owen. 2014. Clathrin adaptors. AP2 controls clathrin polymerization with a membrane-activated switch. Science. 345:459–463.

22. Kirchhausen, T., D. Owen, and S.C. Harrison. 2014. Molecular structure, function, and dynamics of clathrin-mediated membrane traffic. Cold Spring Harb Perspect Biol. 6:a016725.

23. Kovtun, O., N. Leneva, Y.S. Bykov, N. Ariotti, R.D. Teasdale, M. Schaffer, B.D. Engel, D.J. Owen, J.A.G. Briggs, and B.M. Collins. 2018. Structure of the membrane-assembled retromer coat determined by cryo-electron tomography. Nature. 561:561–564.

24. Kremer, J.R., D.N. Mastronarde, and J.R. McIntosh. 1996. Computer visualization of three-dimensional image data using IMOD. J Struct Biol. 116:71–76.

25. Liebschner, D., P.V. Afonine, M.L. Baker, G. Bunkoczi, V.B. Chen, T.I. Croll, B. Hintze, L.W. Hung, S. Jain, A.J. McCoy, N.W. Moriarty, R.D. Oeffner, B.K. Poon, M.G. Prisant, R.J. Read, J.S. Richardson, D.C. Richardson, M.D. Sammito, O.V. Sobolev, D.H. Stockwell, T.C. Terwilliger, A.G. Urzhumtsev, L.L. Videau, C.J. Williams, and P.D. Adams. 2019. Macromolecular structure determination using X-rays, neutrons and electrons: recent developments in Phenix. Acta Crystallogr D Struct Biol. 75:861–877.

26. Mettlen, M., P.H. Chen, S. Srinivasan, G. Danuser, and S.L. Schmid. 2018. Regulation of Clathrin-Mediated Endocytosis. Annu Rev Biochem. 87:871–896.

27. Mettlen, M., and G. Danuser. 2014. Imaging and modeling the dynamics of clathrin-mediated endocytosis. Cold Spring Harb Perspect Biol. 6:a017038.

28. Moore, B.L., L.A. Kelley, J. Barber, J.W. Murray, and J.T. MacDonald. 2013. High-quality protein backbone reconstruction from alpha carbons using Gaussian mixture models. J Comput Chem. 34:1881–1889.

29. Morris, K.L., C.Z. Buffalo, C.M. Sturzel, E. Heusinger, F. Kirchhoff, X. Ren, and J.H. Hurley. 2018. HIV-1 Nefs Are Cargo-Sensitive AP-1 Trimerization Switches in Tetherin Downregulation. Cell. 174:659–671 e614.

30. Morris, K.L., J.R. Jones, M. Halebian, S. Wu, M. Baker, J.P. Armache, A. Avila Ibarra, R.B. Sessions, A.D. Cameron, Y. Cheng, and C.J. Smith. 2019. Cryo-EM of multiple cage architectures reveals a universal mode of clathrin self-assembly. Nat Struct Mol Biol. 26:890–898.

31. Motley, A., N.A. Bright, M.N. Seaman, and M.S. Robinson. 2003. Clathrin-mediated endocytosis in AP-2-depleted cells. J Cell Biol. 162:909–918.

32. Muenzner, J., L.M. Traub, B.T. Kelly, and S.C. Graham. 2017. Cellular and viral peptides bind multiple sites on the N-terminal domain of clathrin. Traffic. 18:44–57.

33. Nickell, S., F. Forster, A. Linaroudis, W.D. Net, F. Beck, R. Hegerl, W. Baumeister, and J.M. Plitzko. 2005. TOM software toolbox: acquisition and analysis for electron tomography. J Struct Biol. 149:227–234.

34. Owen, D.J., Y. Vallis, B.M. Pearse, H.T. McMahon, and P.R. Evans. 2000. The structure and function of the beta 2-adaptin appendage domain. EMBO J. 19:4216–4227.

35. Pettersen, E.F., T.D. Goddard, C.C. Huang, G.S. Couch, D.M. Greenblatt, E.C. Meng, and T.E. Ferrin. 2004. UCSF Chimera--a visualization system for exploratory research and analysis. J Comput Chem. 25:1605–1612.

36. Punjani, A., J.L. Rubinstein, D.J. Fleet, and M.A. Brubaker. 2017. cryoSPARC: algorithms for rapid unsupervised cryo-EM structure determination. Nat Methods. 14:290–296.

37. Qu, K., B. Glass, M. Dolezal, F.K.M. Schur, B. Murciano, A. Rein, M. Rumlova, T. Ruml, H.G. Krausslich, and J.A.G. Briggs. 2018. Structure and architecture of immature and mature murine leukemia virus capsids. Proc Natl Acad Sci U S A. 115:E11751–E11760.

38. Ren, X., G.G. Farias, B.J. Canagarajah, J.S. Bonifacino, and J.H. Hurley. 2013. Structural basis for recruitment and activation of the AP-1 clathrin adaptor complex by Arf1. Cell. 152:755–767.

39. Sali, A., and T.L. Blundell. 1993. Comparative protein modelling by satisfaction of spatial restraints. J Mol Biol. 234:779–815.

40. Schmid, E.M., M.G. Ford, A. Burtey, G.J. Praefcke, S.Y. Peak-Chew, I.G. Mills, A. Benmerah, and H.T. McMahon. 2006. Role of the AP2 beta-appendage hub in recruiting partners for clathrin-coated vesicle assembly. PLoS Biol. 4:e262.

41. Schmid, S.L., A. Sorkin, and M. Zerial. 2014. Endocytosis: Past, present, and future. Cold Spring Harb Perspect Biol. 6:a022509.

42. Schur, F.K., M. Obr, W.J. Hagen, W. Wan, A.J. Jakobi, J.M. Kirkpatrick, C. Sachse, H.G. Krausslich, and J.A. Briggs. 2016. An atomic model of HIV-1 capsid-SP1 reveals structures regulating assembly and maturation. Science. 353:506–508.

43. Shen, Q.T., X. Ren, R. Zhang, I.H. Lee, and J.H. Hurley. 2015. HIV-1 Nef hijacks clathrin coats by stabilizing AP-1:Arf1 polygons. Science. 350:aac5137.

44. Smith, C.J., T.R. Dafforn, H. Kent, C.A. Sims, K. Khubchandani-Aswani, L. Zhang, H.R. Saibil, and B.M. Pearse. 2004. Location of auxilin within a clathrin cage. J Mol Biol. 336:461–471.

45. Taylor, M.J., D. Perrais, and C.J. Merrifield. 2011. A high precision survey of the molecular dynamics of mammalian clathrin-mediated endocytosis. PLoS Biol. 9:e1000604.

46. Tegunov, D., and P. Cramer. 2019. Real-time cryo-electron microscopy data preprocessing with Warp. Nat Methods. 16:1146–1152.

47. Trabuco, L.G., E. Villa, K. Mitra, J. Frank, and K. Schulten. 2008. Flexible fitting of atomic structures into electron microscopy maps using molecular dynamics. Structure. 16:673–683.

48. Turonova, B., F.K.M. Schur, W. Wan, and J.A.G. Briggs. 2017. Efficient 3D-CTF correction for cryo-electron tomography using NovaCTF improves subtomogram averaging resolution to 3.4A. J Struct Biol. 199:187–195.

49. Umasankar, P.K., L. Ma, J.R. Thieman, A. Jha, B. Doray, S.C. Watkins, and L.M. Traub. 2014. A clathrin coat assembly role for the muniscin protein central linker revealed by TALEN-mediated gene editing. Elife. 3.

50. Wan, W., L. Kolesnikova, M. Clarke, A. Koehler, T. Noda, S. Becker, and J.A.G. Briggs. 2017. Structure and assembly of the Ebola virus nucleocapsid. Nature. 551:394–397.

51. Willox, A.K., and S.J. Royle. 2012. Functional analysis of interaction sites on the N-terminal domain of clathrin heavy chain. Traffic. 13:70–81.

52. Wrobel, A.G., Z. Kadlecova, J. Kamenicky, J.C. Yang, T. Herrmann, B.T. Kelly, A.J. McCoy, P.R. Evans, S. Martin, S. Muller, F. Sroubek, D. Neuhaus, S. Honing, and D.J. Owen. 2019. Temporal Ordering in Endocytic Clathrin-Coated Vesicle Formation via AP2 Phosphorylation. Dev Cell. 50:494–508 e411.

53. Xing, Y., T. Bocking, M. Wolf, N. Grigorieff, T. Kirchhausen, and S.C. Harrison. 2010. Structure of clathrin coat with bound Hsc70 and auxilin: mechanism of Hsc70-facilitated disassembly. EMBO J. 29:655–665.

54. Zivanov, J., T. Nakane, B.O. Forsberg, D. Kimanius, W.J. Hagen, E. Lindahl, and S.H. Scheres. 2018. New tools for automated high-resolution cryo-EM structure determination in RELION-3. Elife. 7.

